# Genome-wide Functional Characterization of Escherichia coli Promoters and Sequence Elements Encoding Their Regulation

**DOI:** 10.1101/2020.01.04.894907

**Authors:** Guillaume Urtecho, Kimberly D. Insigne, Arielle D. Tripp, Marcia S. Brinck, Nathan B. Lubock, Christopher Acree, Hwangbeom Kim, Tracey Chan, Sriram Kosuri

## Abstract

Despite decades of intense genetic, biochemical, and evolutionary characterizations of bacterial promoters, we lack the ability to identify or predict transcriptional activities of promoters using primary sequence. Even in simple, well-characterized organisms such as *E. coli* there is little agreement on the number, location, and strength of promoters. We use a genomically-encoded massively parallel reporter assay to perform the first full characterization of autonomous promoter activity across the *E. coli* genome. We measure promoter activity of >300,000 sequences spanning the entire genome and map 2,228 promoters active in rich media. Surprisingly, 944 of these promoters were found within intragenic sequences and are associated with conciliatory sequence adaptations by both the protein-coding regions and overlapping RNAP binding sites. Furthermore, we perform a scanning mutagenesis of 2,057 promoters to uncover sequence elements regulating promoter activity, revealing 3,317 novel regulatory elements. Finally, we show that despite these large datasets and modern machine learning algorithms, predicting endogenous promoter activity from primary sequence is still challenging.

## Introduction

In 1961, François Jacob and Jacques Monod outlined the concept of the bacterial promoter derived from an accumulation of genetic and biochemical studies of metabolic regulation in *Escherichia coli*^1^. Bacterial promoters have since become a foundation for understanding molecular biology and gene regulation, with countless studies probing their genetic, evolutionary, structural, thermodynamic and kinetic properties^2–5^. Several model promoters such as the *lac*, *trp*, and phage promoters have been the subject of in-depth mechanistic studies for how RNA polymerase (RNAP) recognizes promoter sequences, as well as the stepwise process to initiate transcription^6–8^. In addition, many transcription factors have been described in similar detail, revealing the processes through which these proteins modulate the behavior of RNAP and activity of the promoter^4,9–11^. The majority of the binding motifs for these transcription factors have been studied at high resolution using modern methods^12–15^. In short, the myriad components that define *E. coli* promoter function have been extensively cataloged and characterized, establishing them as one of the most well-understood systems in molecular biology.

Despite this extensive knowledge, we still cannot answer many simple and fundamental questions about *E. coli* promoters. For example, how many active promoters exist in *E. coli* at a given growth condition? To what extent is promoter regulation responsible for protein level remodeling during environmental changes? Given a sequence, can we predict whether a promoter is encoded within it as well as its strength and/or regulation? Answers to these questions remain difficult for many reasons. Although the consensus sequences for RNAP recognition motifs have been known for decades, a simple search of the genome based on these motifs yields many false positives. In fact, within a region, there are often sequences closer to the RNAP recognition motifs than the actual functional promoter^16,17^. Experimental efforts to identify promoters using 5’ RNA-Seq have found tens of thousands of putative transcription start sites (TSSs) that presumably mark sites with functional promoter activity, however, there is little overlap between studies^18,19^. Furthermore, although many *E. coli* promoters have been verified with strong biochemical evidence^20^, identifying the cis-regulatory elements responsible for their activity is challenging. As a consequence, roughly two-thirds of the 2,565 reported *E. coli* operons do not contain any transcription factor binding site annotations^20,21^. Finally, aside from a handful of thoroughly-studied promoter sequences, we are still unable to quantitatively predict the activity or behavior of promoters in the context of sequence perturbations such as moving, mutating, or removing transcription factor binding sites.

There are several confounding factors which make it difficult to accurately gauge if a sequence can confer promoter activity. First, recent work has shown that promoter activity varies depending on location in the genome due to factors such as variance in chromosomal copy number^22–24^, the distribution of transcription factors within a cell^25,26^, and chromatin accessibility^27–29^ masking the effects of cis-regulatory elements. Efforts to normalize these effects have utilized reporters on high copy number plasmids that can saturate endogenous transcriptional machinery^30^. Second, inferring promoter strength from endogenous transcript production is problematic because these transcripts often contain sequences that alter their processing and stability independent of the promoter sequence^31,32^. Third, multiple promoters within close proximity, whether co-directional or opposing, can affect each other’s strength and resulting transcription through mechanisms such as RNAP collisions and antisense RNA^33–36^. Finally, not all sequences that initiate RNA transcription are capable of producing mature and translatable RNA^37^.

Here we investigated promoter regulation in *E. coli* using a massively-parallel reporter assay (MPRA) designed to isolate promoter activity from other confounding factors influencing genetic regulation^38^. We measured promoter activity at 17,189 reported TSSs and found that a majority are not autonomous promoter sequences capable of gene transcription. We then measured promoter activity of 321,123 sheared genomic fragments spanning both strands of the *E. coli* genome (8.5x coverage) to elucidate the promoter landscape in rich and minimal media. We then systematically tiled these regions to precisely map promoter boundaries, revealing many regions with multiple promoters in close proximity, as well as many antisense promoters within genes that shape both codon usage and transcription levels. To characterize sequence motifs encoding promoter activity, we performed systematic mutagenesis of 2,057 active promoters and identified cis-regulatory elements affecting promoter activity. With this approach, we characterized the regulatory effects of 568 transcription factor binding sites reported by RegulonDB as well as 2,583 novel sites, thereby providing functionally annotated profiles for promoters driving expression in rich LB media for 1,158 of the 2,565 operons in *E. coli*. Lastly, we trained several machine learning models on these datasets to better understand the features that may be used to classify *E. coli* promoter sequences and quantitatively predict promoter function from sequence.

## Results

### Functional characterization of 17,635 previously reported *E. coli* promoters reveals many are transcriptionally inactive

We first sought to validate promoters and TSSs identified by several genome-wide studies. We assembled previously reported TSSs from three sources: the RegulonDB *E. coli* database^20^ (8,486 TSSs), a directional RNA-Seq study by Wanner et. al^18^ (2,123 TSSs), and a RNA-Seq study by Thomason et. al^19^ (14,868 TSSs). These three sources identify 23,798 unique TSSs active during log-phase growth in rich media with little agreement regarding the location of TSSs between studies and only 93 exact matches shared between all three (**Figure 1A**). Even when we collapsed clusters of TSSs within 20 bp of each other to the most upstream TSS to minimize redundancy, 17,635 unique TSSs remained. These TSSs are likely some combination of true promoters and false positives due to RNA processing, transcriptional noise, or experimental and computational artifacts.

**Figure 1.**
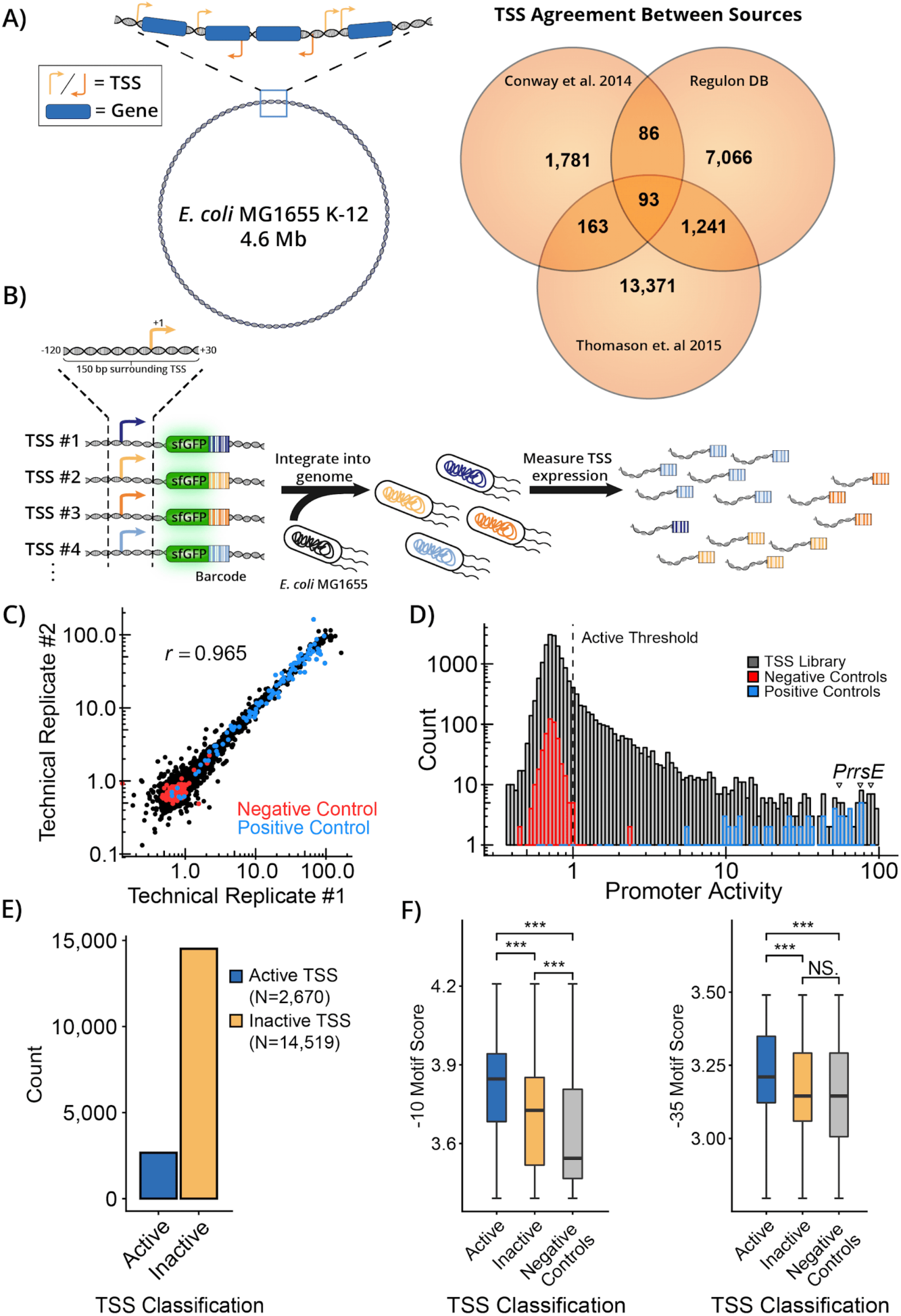
Functional characterization of 17,635 previously reported *E. coli* promoters. **A)** Three sources of genome-wide promoter predictions show little agreement in the reported TSSs at the single-nucleotide level. **B)** We synthesized oligos overlapping the −120 to +30 bp context of 17,635 reported TSSs and integrated construct into a fixed genomic landing pad. Measuring barcode expression using RNA-Seq captures quantitative measurements of transcriptional activity for individual TSSs. **C)** MPRA results are highly replicable across technical replicates (*r* = 0.965, *p* < 2.2 x 10^-16^). **D)** The TSS library measurements span over 100-fold with negative controls exhibiting low levels of expression and positive controls spanning the entire dynamic range. **E)** A majority of tested TSSs are inactive in LB. **F)** Active and inactive TSSs have significantly different mean PWM scores for −10 and −35 σ70 motifs (Wilcoxon rank-sum test, “***” =< 0.001).

To see if these TSS regions could drive gene expression of a transcriptional reporter, we used a genomic MPRA we developed^38^ to quantitatively measure the autonomous promoter activity of 17,635 TSSs (**Figure 1B**). This system allows for single integration of large reporter libraries into a defined locus. The promoter activity reporter is insulated by multiple transcriptional terminators and the reporter transcript contains a RiboJ ribozyme sequence upstream of the RBS that standardizes the transcript produced. For each TSS, we synthesized oligonucleotides spanning 120 bp upstream to 30 bp downstream of the TSS, which encodes the majority of promoter activity driving expression at a given TSS^39^. We included 96 well-characterized promoters from the BioBricks registry ^40^ designed to span a wide range of expression levels to serve as positive controls. We also included 500 negative controls that were selected 150 bp sequences from the *E. coli* genome. Our criteria for selecting these sequences was that they are more than 200 bp from any TSS reported in the aforementioned studies. We engineered these 18,222 unique sequences to express a uniquely barcoded sfGFP transcript and subsequently integrated this pooled library of reporter constructs into the *nth-ydgR* intergenic locus within the *E. coli* chromosomal terminus using a recombination-mediated cassette exchange system^41^. We determined promoter activity levels by performing targeted amplicon sequencing of the barcoded sfGFP transcripts to quantify RNA-seq levels of each barcode normalized to their DNA-seq abundances, and precisely measured expression for 97.5% (17,767/18,222) of TSSs in this library (**Figure 1C**) with an average of 69.5 barcodes measured per library member (**Figure S1A**). Expression measurements were consistent between replicates which were separately barcoded, cloned, and quantified (R^2^ = 0.919, *p* < 2.2 x 10^-16^) (**Figure S1B**). To call a TSS active we set a threshold of at least greater than two standard deviations above the median of the negative control distribution and normalized the data such that the threshold value was set to 1 (**Figure 1D**). Among the 17,635 original TSSs, we confidently quantified 17,189 (97.4%) and identified 2,670 exhibiting expression levels above our experimentally determined threshold (**Figure 1E**). Notably, this number of active promoters is more consistent with the number of operons identified using long-read sequencing to characterize full-length *E. coli* transcripts^42,43^. Amongst these 2,670 confirmed promoters, we recovered expression data for many well-known promoters and three of the strongest corresponded to the 16S and 23S polycistronic operon, the most highly expressed operon in the *E. coli* genome^44^.

To confirm whether our set of negative controls were truly depleted of promoter activity, we tested a set of 936 completely random 150 bp nucleotide sequences and compared the expression levels to our negative controls (**Figure S2**). Despite overall low mean levels of expression (Random sequences: 0.115, Negative controls: 0.036), 2.35% of random promoters drove expression higher than our negative threshold whereas only 0.851% of negative controls exceeded this threshold. A recent study found that 4/40 (10%) random 103 bp sequences exhibited promoter activity^45^ and suggests the frequency of promoter-like activity in overall sequence space is seemingly very high. These results demonstrate that the negative controls used in our assay are depleted in promoter activity, even compared to completely random sequences, and implies that there is negative selection for spurious promoter activity across certain regions of the *E. coli* genome.

### Chromosomal-position specific effects are consistent across diverse promoter sequences

Several recent studies have shown that promoter expression levels can be highly variable between genomic locations^25,27,28^. However, these studies have primarily focused on individual promoters in multiple locations, leaving uncertainty regarding whether these effects are promoter-specific or represent a more widespread phenomenon impacting any promoter at a given position. To study these chromosomal position effects across a wide range of promoters, we integrated the entire TSS promoter library in both left and right chromosomal midreplichores and compared expression measurements between these positions and the *E. coli* chromosomal terminus (**Figure S1C**). Promoter measurements remained highly consistent between locations, although the two midreplichore positions exhibited slightly higher concordance with each other (*r* = 0.97, *p* < 2.2 x 10^-16^), than either midreplichore to the terminus (*r* = 0.95, *p* < 2.2 x 10^-16^). Positive control sequences, which do not contain regulatory elements in addition to the RNAP binding sites, were highly correlated between all locations. We conclude that overall, diverse promoters exhibit similar relative expression levels across genome-positions, although the absolute expression may vary.

### Inactive TSS-associated promoters are enriched for −10 but not −35 σ70 binding motifs

A majority of *E. coli* promoters are regulated by the housekeeping sigma factor σ70^46^, and thus we expected that active promoters would be enriched for the canonical σ70 motifs. Promoters of the σ70 family are well known for containing two hexamer motifs, the −10 and −35 motifs, which recruit RNAP and are named after their position relative to the TSS. We used a σ70 position-weight matrix (PWM)^16^ to analyze whether active TSS promoters were enriched for these motifs. Although both active and inactive TSS-associated promoters were enriched for the canonical −10 motif compared to our negative controls (active: *p* < 2.2 x 10^-16^, inactive: *p* = 6.2 x 10^-8^), we found the −35 scores of inactive promoters were generally no greater than negative controls (*p* = 0.33) (**Figure 1F**). Conversely, active TSS-associated promoters contained significantly higher −35 scores than negative controls (*p* = 1.4 x 10^-8^) or inactive TSS-associated promoters (*p* < 2.2 x 10^-16^). We conclude that inactive TSS-associated promoters lack −35 elements but may become active in growth conditions where additional transcription factors mobilize and facilitate RNAP positioning in the absence of a −35 motif.

### Genome-wide Identification of *E. coli* promoters

Despite functionally screening 17,635 previously implicated TSS regions, we encountered instances where essential operon promoters remained unidentified, suggesting that there were still undiscovered promoters within the genome. For instance, despite screening several reported TSS regions upstream of the essential *yrbA-murA* operon, none exhibited expression greater than our activity threshold. To comprehensively detect all promoters, we cloned, barcoded, and measured the transcriptional activity in LB of 321,123 sheared genomic fragments ranging between 200 and 300 bp (median = 244 bp), providing ∼8.5x coverage per strand of the *E. coli* genome (**Figure 2A, Figure S3A, Figure S3B**). We averaged the expression of fragments overlapping each nucleotide position to achieve highly replicable values of strand-specific promoter activity at single-nucleotide resolution (**Figure S3C**). This data may be viewed using our online tool (https://ecolipromoterdb.com), revealing defined regions of promoter activity across the entire *E. coli* genome (**Figure 2B**). We classify candidate promoter regions as contiguous regions of at least 60 bp with promoter activity measurements higher than an empirically derived threshold. This threshold was established to maximize recall of previously identified active TSSs while minimizing the inclusion of inactive TSSs.

**Figure 2.**
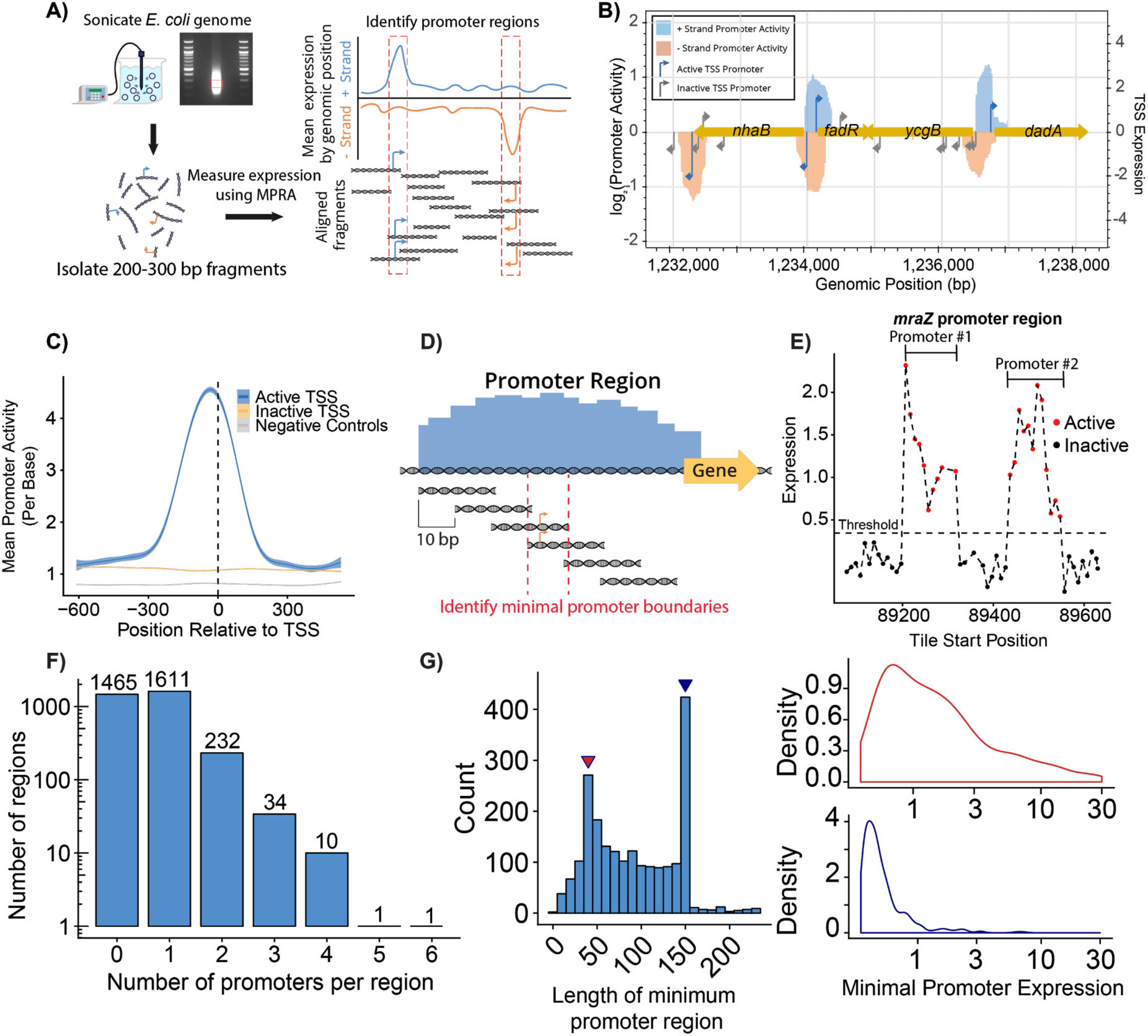
Genome-wide Identification of *E. coli* promoters. **A)** 321,123 sheared genomic fragments were screened using the same MPRA platform. The fragments were 200 to 300 bp in size giving an average 8.5x coverage across each strand of the *E. coli* genome. Promoter activity of each fragment was measured and averaged at each position to recover nucleotide-specific expression. **B)** We created a website to showcase the *E. coli* promoter landscape (https://ecolipromoterdb.com/). This section of the genome displayed in this figure contains five candidate promoter regions that appear within intergenic regions. **C)** Meta-analysis of mean promoter activity at experimentally validated active TSSs, inactive TSSs, and negative controls. **D)** Oligo tiling library identifies promoters within candidate promoter regions. We synthesized 150 bp oligos tiling all promoter regions identified in rich media at 10 bp intervals. We then determine minimal promoter boundaries by identifying the overlap of transcriptionally active tiles. **E)** Oligo tile expression across the *mraZ* promoter shows two distinct promoters. Positions are defined according to the right-most genomic position of each 150 bp oligo. Dashed line indicates the threshold for active oligo tiles **F)** Distribution of the number of promoters per promoter region shows many regions contain multiple promoters. **G)** Left: Distribution of the lengths of the minimal promoter boundaries shows enrichment for σ70-promoter sized regions (40 bp). Right: 40 bp minimal promoters (red) span a wide range of expression whereas 150 bp promoters are typically weak (blue).

With the chosen threshold, we found 3,477 candidate promoter regions in LB that overlapped 2,293/2,670 (85.8%) active TSSs identified in LB, 3,193/14,493 (22.0%) inactive TSSs, and 47/482 (9.75%) negative controls. Active TSSs not overlapping a candidate promoter region generally exhibited weak activity, which may indicate that greater sensitivity is achieved through testing of oligo-array synthesized regions (**Figure S3D**). In many cases, candidate promoter regions overlapped multiple TSS-associated promoters, both active and inactive, indicating the potential for multiple promoters within individual regions (**Figure 2B)**. Overall, we detected strong promoter activity at active TSSs with little promoter activity at inactive TSS promoters, demonstrating agreement between these independent methods for capturing genome-wide promoter activity (**Figure 2C**).

### Fine mapping of *E. coli* promoters within transcriptionally active regions

Our survey of genomic fragments identified candidate regions of promoter activity that were well above the expected size of typical promoters (**Figure S3E**)^39^. To determine if these candidate regions contained multiple promoters, we constructed a library of 48,379 150 bp oligos that tiled the entire lengths of the 3,477 promoter regions identified in LB at 10 bp intervals (**Figure 2D**). For candidate promoter regions under 150 bp, we synthesized a single oligo encoding the region without including additional surrounding sequence context. We recovered highly replicable data for 45,201(93.4%) of these variants with an average of 8 barcodes per variant (**Figure S3F, S3G**). Using this approach, we could precisely pinpoint the boundaries of promoters by observing the specific locations along the promoter region where tiled oligos exhibited changes in expression levels. (**Figure 2E**). This analysis revealed that 1,889 of the previously identified promoter regions contained one or more discrete promoters, including 278 regions containing multiple promoters (**Figure 2F)**. Notably, the number of promoters within a given region correlated with the size of the candidate region (**Figure S3H**) but not necessarily the overall promoter activity of the region (**Figure S3I**). In 1,465 candidate regions, no promoters were detected. These regions typically measured under 150 bp in length, raising the possibility of being false positives or potentially requiring additional transcription factors beyond the scope of the 150 bp regions assessed. Altogether, this approach identified 2,228 distinct promoters active in LB. Furthermore, by determining the overlap of all active oligos tiling a promoter, we were able to infer the minimal sequence necessary for each promoter. When comparing the sizes of the minimal sequence necessary for promoter activity, we observed an enrichment for sequences of approximately 40 bp, which is a typical size for σ70 promoters^47–49^ (**Figure 2G**). We also observed an enrichment for 150 bp minimal promoter regions, although these were generally weak indicating that our resolution is limited when tiling weaker promoters. Overall, we were able to precisely map boundaries for 2,228 promoters active in LB. Considering non-overlapping active promoters identified during our TSS screen, we find 2,859 distinct promoters. Amongst these promoters, we have identified promoters regulating 99 out of 100 randomly sampled essential genes including the promoter for the essential *yrbA-murA* operon which was missed in the TSS screen (**Supplementary Table 1**). The missing promoter was for the *yjeE* gene, which exhibits an atypical operon structure, wherein the first gene in the operon overlaps a gene encoded in the opposite direction. Furthermore, we detected promoter activity in regions 100 bp upstream of 24 of 38 recently described small open reading frames (smORFs) identified by ribosome profiling ^50^, indicating that these proteins may be transcriptionally-regulated independently of larger neighboring genes (**Figure S4**).

### Intragenic promoters are widespread, often found in the antisense orientation, and alter transcript levels and codon usage of the genes they are within

While promoters are commonly thought of as gene regulatory sequences upstream of transcribed genes, they can also be found within genes and oriented to transcribe genes in the antisense direction. We thus sought to explore these atypical promoters and their consequences on the *E. coli* genome and transcriptome. Many studies have found pervasive antisense transcription in prokaryotes ^51–54^, though there is controversy over the functional relevance and whether they are just due to a noisy transcriptional apparatus^55^. At the same time, it has been functionally shown that antisense promoters can alter a sense gene’s transcription, translation, and steady-state message levels^35,56^. Amongst the 2,228 promoters we precisely mapped, 1,131 were primarily encoded within intergenic regions while 944 were found to fully or mostly overlap intragenic regions (**Figure 3A, Figure S5A**). Notably, intragenic promoters exhibited a higher prevalence within single-gene operons compared to individual genes within polycistronic operons (p = 1.05 x 10^-9^, df = 1, Chi-squared Test). Although intergenic promoters were predominantly positioned in the sense orientation relative to the nearest downstream gene, 300 of the 944 intragenic promoters were positioned antisense relative to the genes they overlapped. Interestingly, intragenic promoter activity had greater correlation when comparing activity between growth mediums, indicating that these regions may be primarily composed of constitutive promoter elements (LB: *r* = 0.648, M9 minimal: *r* = 0.787, *p* > 1 x 10^-16^, Wilcoxon rank-sum test, **Figures S5B-C**).

**Figure 3.**
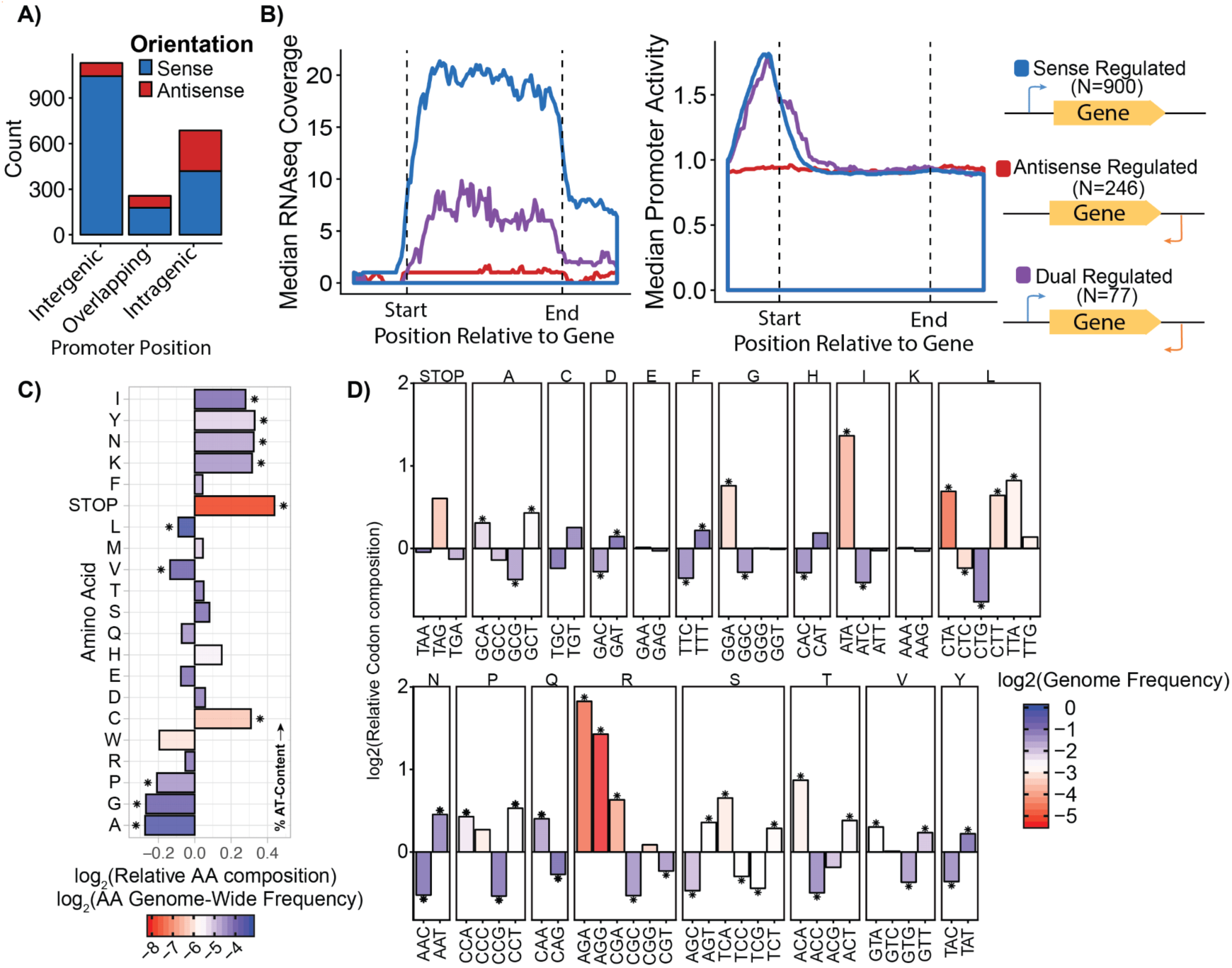
Intragenic promoters are widespread, often found in the antisense orientation, and alter transcript levels and codon usage of the genes they are within. **A)** Orientation and positioning of identified promoters reveals many promoters are intragenic and antisense. **B)** Antisense promoters suppress gene expression genome-wide. Left: Meta-gene analysis of the median RNA-Seq coverage across all sense, antisense, and dual-regulated genes. Right: Meta-gene analysis of sense promoter activity at sense, antisense, and dual regulated genes. **C)** Intragenic promoters are enriched for specific amino acids relative to whole genome amino acid frequencies (Chi-squared test, “*” = *p* < 0.05). Amino acids are arranged by mean AT-content of all corresponding codons. **D)** Specific, often rare, codons are enriched in intragenic promoters. Codon bias within intragenic promoters relative to whole genome. Bars are colored by the relative genome-wide usage compared to other synonymous codons (Chi-squared test, “*” = *p* < 0.05).

Given that we have determined the locations of the antisense promoters driving transcription, we evaluated the genome-wide consequences of antisense promoters on the transcriptome. We performed RNA-Seq on *E. coli* MG1655 grown in LB and compared the transcript coverage of all genes with sense promoters, antisense promoters, and both sense and antisense promoters. We found that overall, genes regulated by both sense and antisense promoters exhibited a two-fold decrease in expression compared to strictly sense-regulated genes (**Figure 3B**). Notably, sense-regulated genes exhibited similar promoter activity on average when compared to genes with both sense and antisense promoters, indicating that the result cannot be attributed solely to stronger promoters in sense-regulated genes. Genes with only antisense promoter activity generally did not exhibit detectable sense transcription.

The significant overlap observed between protein-coding and promoter sequences is interesting given the sequence specificity necessary to encode these distinct functions. Therefore we sought to investigate how sequences navigate this constraint to accommodate diverse activities. After comparing the amino acid composition within intragenic promoters, we observed a significant enrichment of stop codons and a preference for amino acids encoded by codons with higher AT nucleotide content (**Figure 3C**). Further inspection revealed specific codons that were preferentially utilized within intragenic promoter regions (**Figure 3D**), with a notable bias observed among arginine codons, showing a strong preference for AGA and AGG codons. The most enriched codons within intragenic promoters were typically rare in the genome, which may indicate a role of preferential codon usage in controlling promoter activity within genes. The connection between rare codons and regulatory roles has been previously observed in the context of N-terminal codon bias, where rare codons influenced expression levels through secondary structure interactions^57^. Moreover, the observed higher percentage of AT-content^58,59^ and rare codons^60^ may further support the notion that intragenic promoters are linked to horizontally-acquired genes.

Next, we investigated how intragenic promoter sequences had adapted to conform to the constraints of protein-coding sequence space. A peculiar feature of promoter sequences in *E. coli*, is the presence of trinucleotides matching stop codons within the canonical −10 and −35 σ70 motifs (−35: 5’-T**TGA**CA, −10: 5’-TA**TAA**T). Therefore, we hypothesized that the reuse of these nucleotide patterns offers another mechanism by which the *E. coli* genome counteracts the spurious evolution of intragenic promoters, thereby explaining their scarcity relative to the ease by which they can evolve^45^. We used a σ70 PWM^16^ to identify the highest-scoring σ70 motifs within intragenic promoters and determined their relative coding frames. Interestingly, we observed a lower frequency of −35 elements in +2 coding frames and the −35 motifs detected at +2 positions exhibited significantly reduced resemblance to the canonical motif (**Figure S6A**). Similarly, −10 motifs were least frequently found in the +1 positions, although −10 motifs at this position did not show lower overall scores (**Figure S6B**). The observed depletion of −35 motifs positioned in the +2 reading frame and −10 motifs in the +1 reading frame is likely due to the fact that the canonical sequences for these motifs would create stop codons within the protein if placed at these positions. This suggests a simple, but effective preventative mechanism against the spurious evolution of intragenic promoters that is inherent to their sequence motifs.

### The *E. coli* promoter landscape is dynamic in response to environmental conditions

It is well understood that bacterial cells respond to environmental conditions through changes in their transcriptional profiles^61^, however, it has not been shown how the global promoter landscape changes to facilitate these cellular transitions. To explore this, we measured promoter activity of our genomic fragment library in exponentially growing cells under glucose minimal media conditions. Compared to LB, cells grown in glucose minimal media do not have access to environmental amino acids and must synthesize these and other essential compounds on their own^62^. We recovered replicable promoter activity measurements for 318,457 genomic fragments in glucose minimal media, spanning the genome with 8.38x coverage (**Figure S7A, Figure S7B**). We identify 3,321 candidate promoter regions in glucose minimal media with an average length of 293 bp (**Figure S7C**). Although 2,466 of these regions overlapped with regions found in LB, we found 960 only found in LB and 1,029 exclusive to M9 **(Figure 4A)**. Many of these condition-dependent promoter regions were weak compared to those identified in both conditions **(Figure S7D)**, nonetheless, each condition revealed distinct strongly activated regions unique to it. The observed low activity of condition-unique promoters is similar to what has been observed in synthetic inducible promoter systems, where tightly-regulated promoters often exhibit reduced expression in induced conditions^63^. To identify the most differentially-expressed promoters in each condition, we extracted regions larger than 60 bp that exhibited greater than two-fold difference in activity between conditions. With this criterion, we found 278 regions upregulated in LB and 644 regions upregulated in glucose minimal media. In glucose minimal media, the greatest increase in promoter activity occurred at *ryhB*, a Fur-regulated gene encoding a small RNA that regulates iron-binding and iron-storing proteins when available iron is limited^64,65^ **(Figure S7E).** In LB, the strongest activated region is positioned to drive expression of the *rbsDACBKR* operon, which is essential for uptake and utilization of extracellular ribose^66^ **(Figure S7E)**.

**Figure 4.**
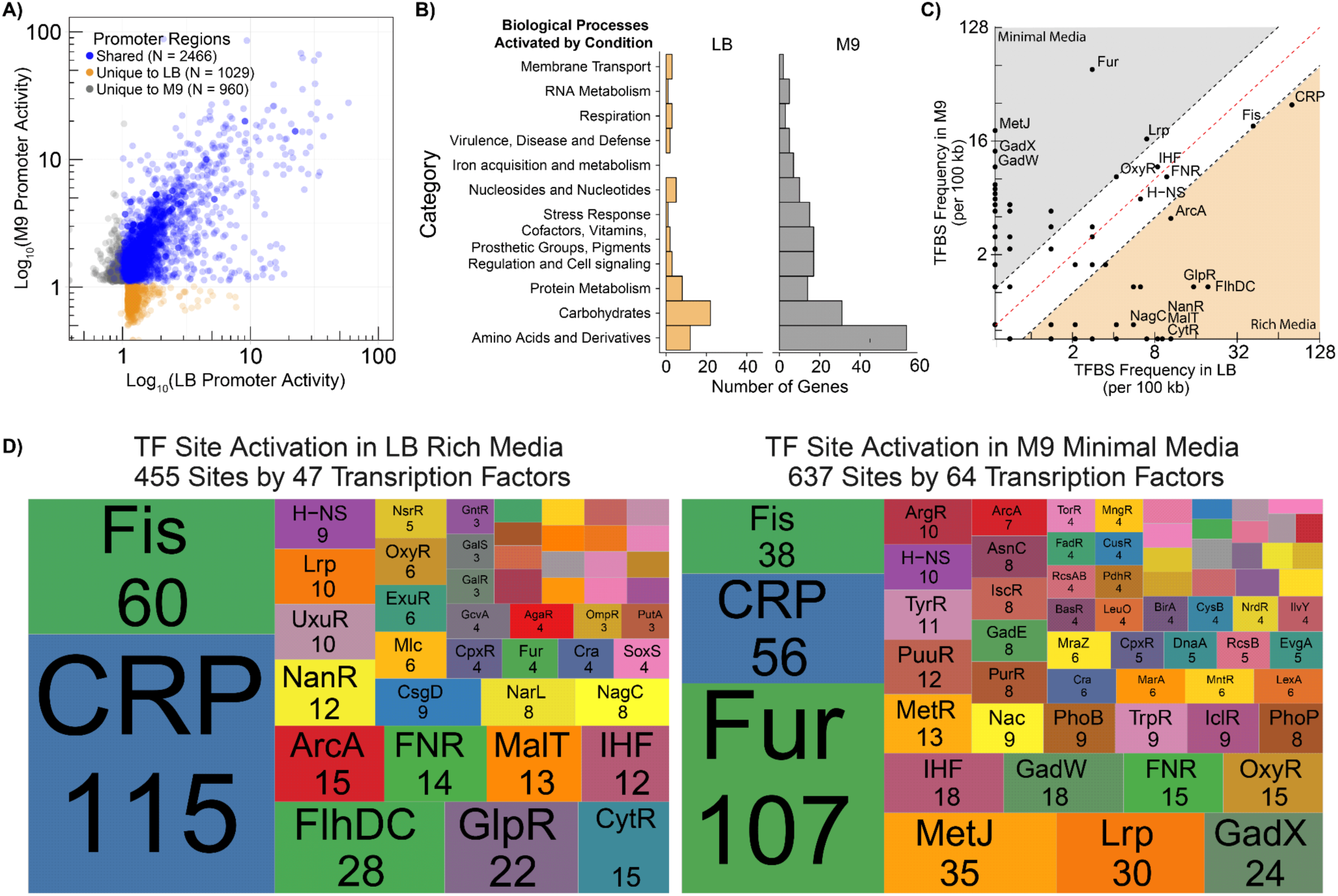
The *E. coli* promoter landscape dynamically responds to environmental conditions. **A)** Shared and unique promoter regions are found between LB and glucose minimal media. **B)** Genes activated by promoters in glucose minimal media are enriched for amino acid-related genes according to RAST subsystem annotations. **C)** Occurrence of reported transcription factor binding sites in promoter regions activated in LB compared to glucose minimal media (M9). Black lines indicate 2-fold enrichment threshold. **D)** Number of binding sites per transcription factor within activated promoter regions. A median of four sites per transcription factors were activated in LB and a median of five sites in M9.

For each condition, we matched activated intergenic and sense promoter regions with the nearest downstream gene and found 159 genes poised for activation in LB and 392 genes poised for activation in glucose minimal media. To see if promoter activation resulted in an increase in expression of these genes, we compared RNA-Seq coverage of the genes with the top 100 strongest promoter activation in each condition (**Figure S8**). In each condition, promoter activation resulted in a concomitant increase in RNA-Seq coverage (LB: *p* = 1.1 x 10^-5^, M9: *p* = 1.9 x 10^-5^, Wilcoxon rank-sum test). To see which cellular responses were being mobilized by remodeling the promoter landscape, we used the RAST annotation engine^67,68^ to assign functional categories to activated genes and identify enriched cellular processes. Genes downstream of promoter regions activated in LB are predominantly associated with carbohydrate utilization whereas genes downstream of promoters activated in glucose minimal media were associated with amino acid utilization (**Figure 4B**). Overall, we find distinct condition-dependent activation of promoter regions leading to changes in gene expression associated with carbohydrate utilization in LB and amino acid utilization in glucose minimal media.

Next, we explored how these changes in the promoter landscape are mediated by transcriptional machinery and evaluated the transcription factor binding site (TFBS) composition of promoter regions activated in each condition. As opposed to traditional transcriptomebased measurements which measure changes in downstream gene expression, this assay identifies upstream regulatory regions that contribute to promoter activity in response to changing conditions. By cross-referencing these activated promoter regions to TFBSs reported by RegulonDB, we identified transcription factors facilitating these changes to the promoter landscape (**Figure 4C**). Upon comparing TFBS content of these regions we found that binding sites for several global transcriptional regulators^69^, including IHF, Lrp, and Fis occurred at similar frequencies between these conditions. Conversely, binding sites for Fur, another global transcription factor, were enriched by roughly 20-fold within regions activated in glucose minimal media compared to regions activated in LB. This transcription factor is essential for maintaining iron homeostasis^70,71^, and is a known regulator of *ryhB,* the most upregulated gene we found in glucose minimal media. Binding sites for CRP were enriched by more than two-fold in regions activated in LB compared to glucose minimal media. This transcription factor is activated in glucose-limited conditions and so would likely not induce promoter activity in glucose minimal media. Overall, we found 455 TFBSs within regions activated in LB and 637 annotations in regions activated in glucose minimal media (**Figure 4D**). In addition to global regulators, we found many TFBSs that appear exclusive to each condition targeting relatively few regulatory targets. Interestingly, the combined contribution of non-global transcription factors activating 10 or fewer sites were responsible for over a third of all activated promoter regions, underscoring the significant involvement of local transcriptional regulators in driving the overall changes to the transcriptome. Transcription factors MetJ, GadX, and GadW were exclusively found in regions activated in glucose minimal media whereas FlhDC, GlpR, and CytR were the most enriched amongst regions activated in LB.

### Mutational scanning of 2,057 *E. coli* promoters identifies regulatory elements controlling transcription

After globally identifying promoter regions in the bacterial genome, we sought to develop an approach to identify sequence motifs regulating these promoters. Recent work has demonstrated a high-resolution saturation mutagenesis approach to identify regulatory motifs within individual uncharacterized promoters^21,72^. Inspired by this work, we implemented a scanning mutagenesis strategy to explore the sequence features that regulate active promoters. For 2,057 active TSS-associated promoters identified in LB, we systematically scrambled individual 10 bp sequences spanning the −120 to +30 positions at five bp intervals (**Figure 5A**). Using this approach, we would expect that disrupting a repressor site would increase expression, whereas disrupting a RNAP or activator site would decrease expression. These scrambled sequences were designed to maximize distance from the original sequence while maintaining nucleotide content, ensuring perturbation of any motifs at each position contributing to transcriptional regulation. In total, we designed a library of 59,653 sequences consisting of 2,057 active TSS-associated promoters, their scrambled variants, and the previously described set of negative and positive controls. We measured promoter activity of this library as before and recovered replicable expression measurements for 52,900/59,653 (89%) of this library in LB, with an average of seven barcodes per variant (**Figure S9A, S9B**). Using this approach, we identify regions that either increased or reduced expression across thousands of promoters in a single assay (**Figure 5B**). These sequences were enriched at the −35 and −10 positions for regions that increased expression, which is expected considering the majority of promoters are σ70 dependent. However, many sequences outside of these −10 and −35 regions were also found to contribute to regulation.

**Figure 5.**
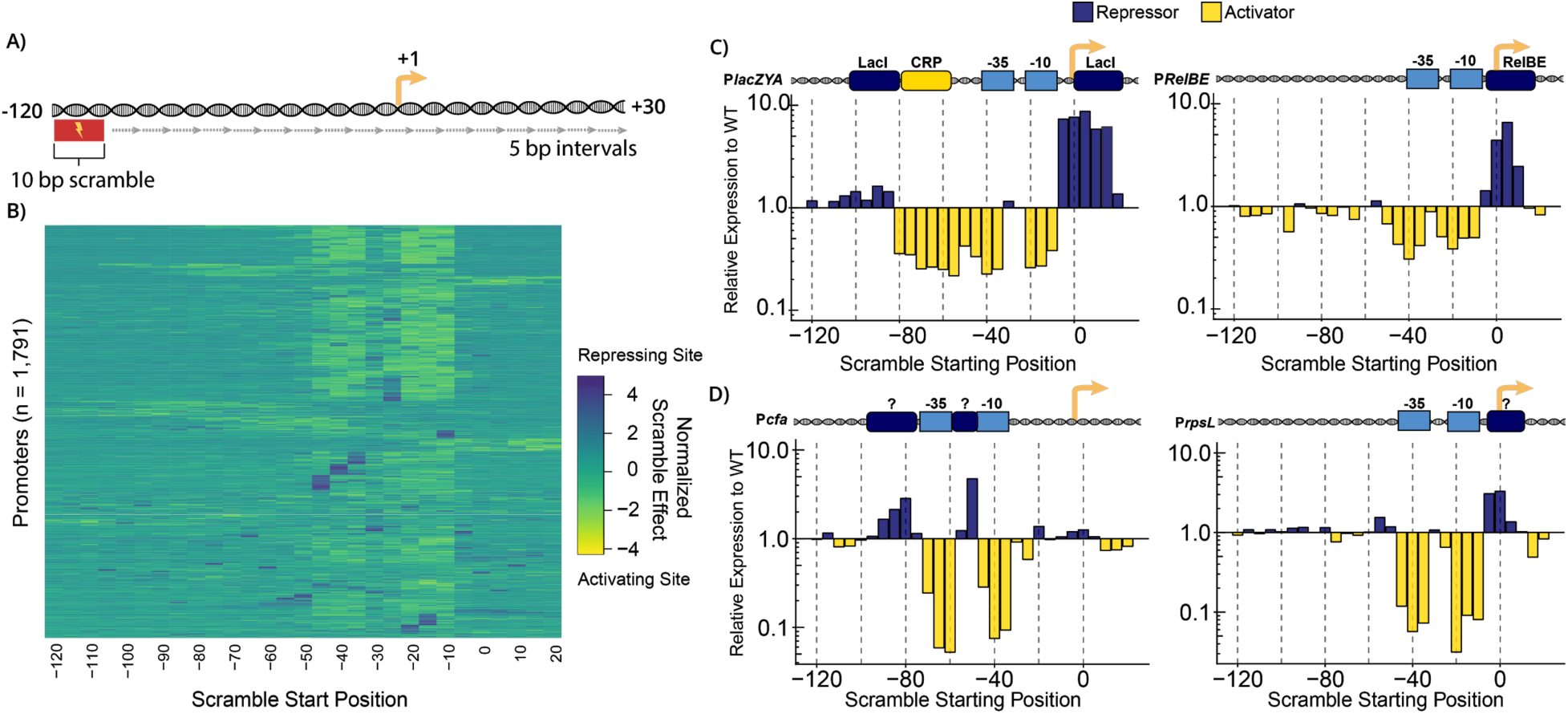
Scanning mutagenesis of 2,057 TSS-associated promoters identifies known and novel regulatory motifs. **A)** Scanning mutagenesis of 2,057 *E. coli* promoters to identify regulatory elements. For each promoter, 10 bp regions were mutated across the full length of the promoter at 5 bp intervals. **B)** Mutating each position across *E. coli* promoters identifies sequences that activate and repress promoter activity. Rows are rearranged using hierarchical clustering and the intensities are normalized within each row. **C)** Scanning mutagenesis of the well-characterized (Left) *lacZYA* and (Right) *relBE* promoters captures known regulatory elements. **D)** Scanning mutagenesis of the newly characterized (Left) *cfa* and (Right) *rpsL* promoters identifies regions encoding regulation within these promoters.

To validate our approach, we first examined the *lacZYA* promoter, a classic gene regulation model whose sequence motifs are well characterized. This promoter is known to contain a variety of regulatory motifs, including twin LacI repressor sites centered at +11 and −82^73^, a CRP activator site centered at −61^74^, and a σ70 RNAP binding site. Our analysis revealed distinct signals corresponding to each of these sites, as well as quantitative measurements for their contribution to expression (**Figure 5C**). Additionally, scanning mutagenesis of the previously characterized *relBE* promoter achieved similar results, identifying a reported RelBE repressor site at the +1 position^75^ as well as −10 and −35 σ70 recognition motifs^75^.

Considering that our approach effectively captures the effects of known binding sites, we proceeded to investigate whether it could also identify regulatory sites within uncharacterized promoters. Although we performed this scanning mutagenesis for 2,057 TSS-associated promoters, here we highlight a few examples to demonstrate the utility of this method (**Figure 5D**). The cyclopropane fatty acyl phospholipid synthase gene, *cfa,* exhibits dynamic expression^76^ and plays a crucial role in cell membrane integrity under acidic conditions ^77^. While there have been several transcription factors implicated in regulation of *cfa*, the motifs responsible for its direct regulation are still unknown. Our approach identified a candidate σ70 promoter regulating this gene with a −10 motif centered 34 nucleotides upstream of the reportedly associated TSS as well as a −35 motif 57 bp upstream, implying that the reported TSS is likely not the primary site for transcription initiation. Additionally, we identified two repressor sites—one located in the spacer region and another upstream of the −35 motif. We also identified novel regulatory regions for an uncharacterized promoter regulating *rpsL*, an essential gene and component of the 30S ribosomal subunit. In this case, we identified a candidate σ70 RNAP binding site with predictably positioned −10 and −35 motifs, as well as an unknown repressor located over the transcription start site. Notably, mutating the repressor site resulted in a threefold increase in promoter expression. Although further experiments^21^ are necessary to identify the transcription factors acting on these promoters, our results provide valuable insights by pinpointing the sequence elements responsible for the regulation of these genes.

### Global identification of 7,293 *E. coli* promoter regulatory motifs

We expanded the scope of our analysis to systematically explore the regulatory motifs amongst all 2,057 promoters tested. We used individual barcode measurements, across four replicates, to find significant differences between the mean expression of the WT and mutated sequences (Student’s t-test with 1% FDR). Among the mutations that significantly altered expression, 1,885 increased expression whereas 5,408 decreased expression (**Figure 6A**). Mutated sites were located throughout promoters and resulted in dramatic changes in expression, some over 100-fold (**Figure S10A**). We observed markedly different distributions for the positions of sequences that increased expression compared to those causing decreased expression (**Figure 6B**). Regions that increased expression were enriched at the −10, −35, and −70 positions, which is consistent with the σ70 RNAP binding motif as well as the typical position of transcriptional activators among class I bacterial promoters^78–80^. Regions that decrease expression were found to localize to the TSS, spacer, and −35, which is consistent with known mechanisms of RNAP occlusion by steric hindrance^80,81^. Alternatively, repressive sites within the spacer could be negatively influencing transcriptional initiation through transcription factor-independent mechanisms ^82^. Furthermore, we found that intergenic promoters contained more regions that altered promoter activity when scrambled compared to intragenic promoters, implying that intragenic promoter sequences contain more compact or fewer regulatory elements (**Figure S10B**).

**Figure 6.**
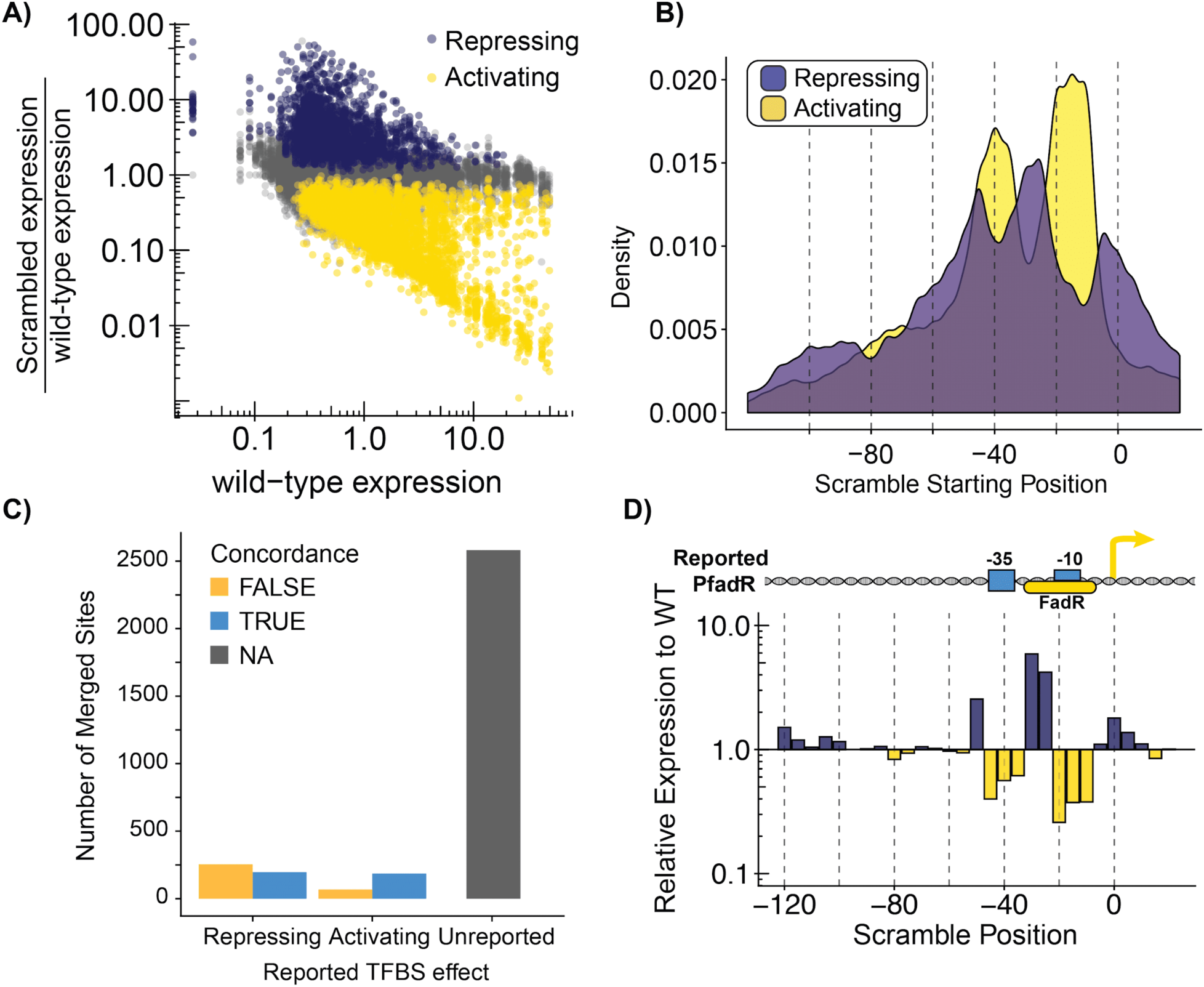
Global identification of 3,317 *E. coli* regulatory motifs by scanning mutagenesis. **A)** We identified scrambled regulatory regions that significantly increase (N = 1,885) or decrease (N=5,408) expression when scrambled relative to the unscrambled promoter. Data are colored by whether the regulatory region activates or represses activity of the promoter. **B)** Activating promoter sequences are enriched at the −10, −35, and −80 positions whereas repressing sequences are enriched at +1, −20, and −50 positions. **C)** Identified regulatory regions overlapping reported TFBS annotations shows mixed concordance with reported effects; 77.8% (2,583/3,317) of identified regulatory regions are unreported by RegulonDB. **D)** Scanning mutagenesis of the *FadR* promoter (bottom) identifies a repressing sequence near the −30 that has been reported to be activating (top).

Next, we cross-referenced these regulatory regions with the extensive collection of putative and experimentally determined regulatory sites reported by RegulonDB^83^. First, for all promoter mutagenesis profiles, we merged adjacent regions found to influence promoter activity, resulting in the identification of 1,414 regions that increase expression and 1,903 regions that decrease expression. Sites were 20 bp on average (indicating they exhibited regulatory impacts across four consecutive 10 bp scramble mutations spaced 5 bp apart) (**Figure S10C**) with effect sizes largely independent of their lengths (**Figure S10D**). Of the 2,453 unique TFBSs reported by RegulonDB, 1,156 overlap with regulatory regions identified by our analysis and 49% (567/1,156) resulted in a significant change in activity of the promoter. The effect we observed after disrupting these reported TFBSs often did not agree with the annotated effect. Our scrambling results agreed with the reported effect for 65% (185/253) of activators and 43% (196/450) of repressors (**Figure 6C**). We presumed the lower concordance with repressors could be due to scrambling mutations disrupting both a repressor and −35 or −10 element, resulting in a decrease in expression which would appear to contradict a reported repressor site. Looking at the distribution of concordance for merged scrambles by position relative to the TSS, we observed a higher proportion of disagreement near the −35 and −10 elements, suggesting overlapping scrambles may be disrupting crucial promoter elements in addition to reported repressor sites (**Figure S10E, S10F**). This may be expected considering that many repressors operate by binding regions proximal to the RNAP binding site. Regardless, we found several examples where the regulatory effects predicted by RegulonDB were contradicted with strong evidence, which may indicate that the effect of the reported annotation is incorrect or that these sites may support multiple transcription factors (**Figure 6D**). Overall, we characterized regulatory sequences in promoters driving expression of 1,158 of the 2,565^83^ operons in *E. coli* as well as many other confirmed promoters. Thus, we conclude that this approach is an efficient and effective method to rapidly characterize regulatory motifs within thousands of experimentally verified promoter regions.

### Predicting promoter activity from sequence remains a challenge

In this study we generated a powerful dataset linking 117,556 unique 150 bp sequences to a quantitative measurement of *in vivo* promoter activity. Using this unique dataset, we evaluated our ability to determine whether a promoter was active or inactive (classification) and the precise level of activity (regression). We trained several machine learning models of varying complexity for both classification and regression. As many sequences are highly similar due to library design and close proximity of previously reported TSSs, we split the data into 75% for training (n = 87,164) and 25% (n = 30,392) for testing according to genomic location, ensuring the two sets contain sequences equidistant to the origin (see Methods). For classification, we determined a threshold independently for each library based on the negative controls. Sequences are considered active if their expression is greater than two standard deviations above the negative median value and inactive if expression falls below this threshold.

We trained several different classifiers to predict whether a given sequence was active or inactive (**Figure 7A**). All classifiers output the predicted probability for each class, rather than directly predicting the class, allowing them to be compared using precision-recall curves. Further details for all models are included in the methods. We trained a simple logistic regression based on four biophysical features known to be associated with promoter strength: max −10 σ70 motif position weight matrix (PWM) score, max σ70 −35 motif PWM score, paired −10 and −35 PWM score (PWMs scanned jointly allowing for, 16, 17, or 18 gap between the −10 and −35), and percent GC content. We trained this model only using variants from the TSS library, which contained the greatest diversity, as the model was unable to converge when trained on the full dataset. For comparison, we trained a gapped k-mer SVM (gkm-SVM) model with word-length 10 and 8 informative columns (L = 10, K = 8) on the same training set, as this model is best suited for sample sizes under 20,000 and observed decreased performance relative to the logistic regression (AUPRC = 0.43, AUPRC = 0.53, respectively). Furthermore, we created a feature set of all 3 to 6-mer frequencies and trained a logistic regression, partial least squares discriminant analysis (PLS-DA), and multi-layer perceptron (MLP). To observe the effects of reducing dimensionality, we additionally trained on only 6-mer frequencies for the MLP and random forest. For the simpler logistic regression and PLS-DA we performed an additional feature selection step based on the performance of a random k-mer. All models performed similarly, with AUPRC ranging from 0.26 to 0.33.

**Figure 7.**
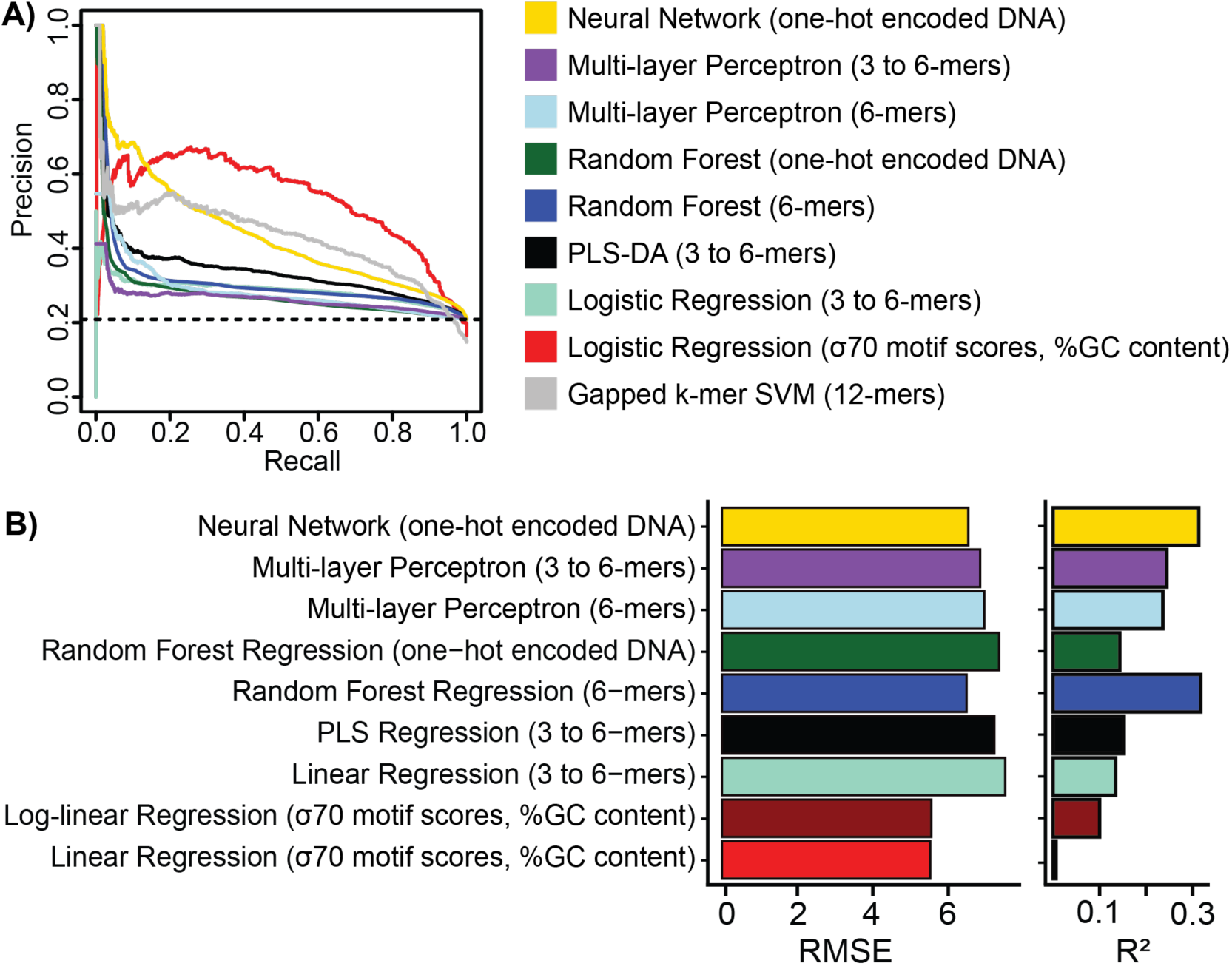
Various machine learning models for promoter activity classification and regression. **A)** Performance of various models to classify promoter sequences. Convolutional neural networks performed best in the lower recall range, while logistic regression based on simple hand-crafted features performs better in the higher recall range. Dashed line represents the expected performance from random prediction using full library. **B)** Performance of regression models to predict a quantitative level of promoter activity. We evaluated performance using both root mean squared error (RMSE) and coefficient of determination (*R*^2^) on the held-out test set. Similar to classification, convolutional neural networks performed the best with the lowest RMSE and highest *R*^2^.

There has been recent work predicting transcriptional regulatory activity from MPRA data using convolutional neural networks (CNNs), which capture intricate sequence features without *a priori* knowledge^84^. Inspired by this work, we trained a CNN using the DragoNN toolkit which is built on top of the keras python package^85^. We performed hyperparameter tuning for a three-layer CNN and achieved an AUPRC = 0.44. Next, we compared the CNN to other machine learning models that require less hyperparameter tuning and are more interpretable. For comparison, we trained a random forest on one-hot encoded DNA, which is not well suited to categorical features, and achieved an AUPRC = 0.27. Furthermore, we trained this model using frequencies of 6-mers and observed a slight increase in performance (AUPRC = 0.31). Overall, the CNN achieved the highest AUPRC, but the logistic regression fit with biophysical features more accurately at higher levels of recall. However, these two models may not be directly comparable, as the logistic regression was trained on only the TSS library rather than the full dataset.

We separately trained all of the models described above, with the exception of gkm-SVM, for the more difficult task of regression (**Figure 7B**). Additionally, we included a linear regression model that fit to the four “mechanistic” features to predict log-transformed expression. We evaluated each model using root mean squared error (RMSE) and *R*^2^ between predicted and observed values for promoter activity. Many models perform similarly to each other, with the CNN achieving the highest R-squared and lowest RMSE (RMSE = 3.12, *R*^2^ = 0.31, *p* < 2.2 x 10^-16^). We observe improvement in the linear regression on log-transformed data compared to linear regression without transformation, suggesting there are non-linear relationships that are presumably captured by more complex models. Random forest on one-hot encoded DNA performs worse than random forest on 6-mer frequencies, in line with the heuristic that random forests are not well suited to categorical features. Overall, the CNN performs best in both classification and regression, although simpler models have some predictive power and have the benefit of faster training times.

## Discussion

More than fifty years have passed since the first conceptions of what bacterial promoters were. Today, *E. coli* promoters are arguably the most well-studied gene regulatory element and yet we cannot seem to agree on basic questions of how many promoters exist, what elements define their function, how constrained they are in sequence space, and how far are we from predicting promoter activity from sequence. Systematic identification and characterization based on transcriptional profiling is confounded by genomic location, RNA processing, stability, and detection differences due to differences in sequences expressed.

Here we attempted to separate promoter activity from other mechanisms of gene regulation to systematically identify promoter locations, strength, and internal structure genome-wide in rich media conditions. We systematically probed previous predictions and combined them with more unbiased approaches to better understand promoter architecture in *E. coli.* Overall, we found 2,859 ≤150bp promoters during log-phase growth in LB, which is consistent with recent estimations by RNA profiling using long-read sequencing technologies^42,43^ and in vitro transcriptional assays^86^. This included many promoters contained within genes, often in the antisense direction, that had large effects on mRNA levels and constrained codon choice within these genes. Despite the ability of our approach to interrogate promoter activity across the entire genome, there are certainly many more condition-specific promoters that remain undiscovered. Moreover, it is likely that we have not identified all active promoters even under the conditions investigated in this study. It is essential to acknowledge that our approach to classifying sequences as promoters is based on an empirically derived threshold. However, this is a simplification as promoters that fall below the threshold could become active due to the influence of other factors, such as message stability^31^ and genomic context^25,27,28^. Taken together, these measurements provide one of the richest datasets on autonomous promoter activity. Our data suggests that all sequences have some propensity to be a promoter, and this propensity is further modulated by other factors such as stability of the message produced or integration locus to ultimately determine mRNA levels. Moreover, the frequency of promoter-like activity in overall sequence space is seemingly very high. This view is consistent with the surprising ease by which promoters evolve from random sequences^45,87,88^.

Our scanning mutagenesis of active TSS-associated promoters identified 3,317 regions with no corresponding TFBS annotation in RegulonDB, revealing that there is a great deal more we can learn about how regulation is encoded in the *E. coli* genome. For regions that overlapped known sites, an appreciable proportion disagreed with the reported effect. There could be several explanations for this disagreement and the discovery of these missing annotations. First, it could be that the predictions of TFBSs in RegulonDB are actually false positives due to promiscuous or nonproductive binding events. This seems plausible considering a recent study of the global transcription factor PhoB, which supports the notion that transcription factors engage in many genomic binding events with apparent non regulatory functions^89^. Second, some transcription factors may possess condition-dependent behavior and the conditions tested in our study do not capture the full scope of their regulatory program. Finally, it is plausible that a portion of the sites we identify represent true functional sites that are missing from current annotation and should be interesting targets for further dissection, such as identifying which transcription factors operate at these motifs. Further studies using high resolution mutagenesis strategies^21^ will be an effective approach to determining which sequences within promoters contribute to regulation and further efforts to predict promoter sequence-function relationships.

To better understand how promoter activity is modulated by sequence, we trained a suite of machine learning models to identify promoter sequences (classification) and predict the precise level of activity (regression). These models varied in complexity, from simple linear regression models based on a handful of known biological features to CNNs trained on raw sequence. Even with the large training set and a wealth of mechanistic information, the performance of these predictive models is limited. There are several possible explanations for why it remains a challenge to classify or predict the activity of *E. coli* promoters. First, it is likely challenging to develop a single generalizable model for all promoters as there are several families of sigma factors with distinct motifs. Therefore, models that are sigma-factor specific may be more tractable. Indeed, recent studies by us and others have leveraged large MPRA datasets characterizing σ70 promoters to develop a variety of statistical and biophysical models that predict expression with surprisingly high accuracy^38,90–92^. These findings suggest that overcoming the challenges associated with promoter activity prediction is plausible with the appropriate training sets and a reasonable scope of study. Second, although the range of our MPRA is quite dynamic, accurate predictive models may require techniques with even greater quantitative resolution, especially in the noise regime of the assay where most observations fall. Finally, we might simply lack the basic models for how sequences define biological functions, such as promoter activity, and thus we are looking in the wrong places for information. Recent efforts to use much larger libraries of random DNA sequences to identify strong promoters may serve as a better starting point to constrain computational models for how sequences affect function^93,94^.

The experimental workflows demonstrated here enable the rapid and iterative exploration of how sequence affects bacterial promoter function. The convergence of DNA synthesis technologies with multiplexed assays for genetic function now allow an individual to routinely design, build and test 10^4^-10^5^ designs on a monthly basis. Such empirical power has no equivalent in other physical systems and has now reached the limits of human experimental design and planning. Thus, understanding bacterial promoters might be one of the best problems to develop and test large-scale design-of-experiment and active learning methodologies to build better predictors and discriminate between different mechanistic models of function.

## Acknowledgements

This work was supported by the National Science Foundation Graduate Research Fellowship 2015210106 and the HHMI Hanna H. Gray Postdoctoral Fellowship (GT15182) to G.U., National Institutes of Health New Innovator Award DP2GM114829 to S.K., Searle Scholars Program (to S.K.), U.S. Department of Energy (DE-FC02-02ER63421 to S.K.), UCLA, and Linda and Fred Wudl. We thank the UCLA BSCRC high throughput sequencing core and Technology Center for Genomics and Bioinformatics for technical assistance; Robert B. Phillips, Reid C. Johnson, and Jeffery H. Miller for thoughtful feedback throughout this project; Matteo Pellegrini for computational advice; Christina P. Burghard for advice on bioinformatics analysis; and all past and present members of the Kosuri lab for technical feedback. Furthermore, we would also like to thank David Gray and Lisa M. Golden for manuscript feedback. We would also like to thank the invaluable resources of RegulonDB and EcoCyc as well as all contributors to these collections. Lastly, we thank the UCLA Molecular Biology Interdepartmental Graduate Program and UCLA Bioinformatics Interdepartmental Graduate Program.

## Author Contributions

G.U., K.D.I., H.K., and S.K. designed the study. G.U., A.D.T., and M.B. developed and performed experimental methods. N.B.L. developed the genomic fragmentation isolation method. G.U., K.D.I., and A.D.T. analyzed, and interpreted data. T.C. developed k-mer based multilayer perceptron for promoter prediction. K.D.I. developed and implemented machine learning approaches for promoter prediction. C.A. and G.U. created the interactive website for data sharing. G.U., K.D.I, and S.K. wrote the manuscript.

## Declaration of Generative AI and AI-assisted technologies in the writing process

During the preparation of this work the author(s) used Chat GPT (GPT-3.5) in order to improve language and clarity. After using this tool/service, the author(s) reviewed and edited the content as needed and take(s) full responsibility for the content of the publication.

## Inclusion and Diversity

One or more of the authors of this paper self-identifies as an underrepresented ethnic minority in their field of research or within their geographical location. One or more of the authors of this paper self-identifies as a gender minority in their field of research. One or more of the authors of this paper received support from a program designed to increase minority representation in their field of research.

## Methods

### Strains

All experiments were performed in the *E. coli* MG1655 background^95^ which carries the following genotype: F-, *λ^-^*, *rph-1* (Yale Coli Genetic Stock Center no. 6300). For the genomically-integrated MPRA, previously reported strains^38^ with engineered landing pads in the right midreplichore (*essQ-cspB* intergenic locus, Addgene no. 110244), chromosomal terminus (*nth-ydgR* intergenic locus, Addgene no. 110245), and left midreplichore (*ybbD-ylbG* intergenic locus, Addgene no. 110243) were used. Briefly, these landing pads encode a fluorescent mCherry reporter as well as chloramphenicol resistance, both of which are flanked by *loxP* sites for recombination-mediated cassette exchange.

### TSS library design

The TSS library incorporates all TSSs from the RegulonDB database^83^ (Version 8.0, genome version U00096.2) and those identified in two recent genome-wide TSS mapping studies^18,19^. Recent work provides evidence that most regulatory motifs fall within 100 bp upstream of the TSS^39^ and the initial transcribed region (+1 to +20) can also influence gene expression. Thus, each TSS was synthesized embedded in its local sequence context −120 to +30 relative to the TSS, capturing a majority of the *cis*-regulatory elements. There were 23,798 unique TSSs across all three sources, many of which were a few base pairs away from each other. We minimized redundancy and collapsed together TSSs within 20 bp and selected the most upstream TSS for our library, yielding 17,635 TSSs for the final synthesized library. Additionally, we included 500 negative controls from the *E. coli* genome that are assumed to have minimal regulatory activity. These were selected from 150 bp regions that are more than 200 bp from a TSS (on either strand), and many fall within coding regions. We included a set of 112 short synthetic positive controls that were previously characterized^40,96^ and span a wide range of expression.

### TSS library barcoding and cloning

The TSS library was synthesized by Twist Biosciences and delivered lyophilized as a 26 pmol pool. The library was resuspended in 100 uL of TE pH 8.0 and 1 uL was amplified for 12 cycles using GU72 and GU116 with NEB Q5 High-Fidelity 2x Master Mix (#M0492L). Unless otherwise stated, all amplifications were performed using this polymerase mixture. This product was then ran on a 2% TAE agarose gel and approximately 200 bp amplicons were extracted using a Zymoclean Gel DNA Recovery Kit (#D4008). For barcoding, 1 ng of this eluate was amplified for 9 cycles using primers GU72 and GU73. Following cleaning using a Zymo Clean and Concentrator Kit (#D40140), the library was digested using NEB’s SbfI-HF and XhoI.

The plasmid backbone, pLibacceptorV2 (Addgene #106250) was digested using SbfI-HF and SalI-HF with the addition of rSAP (NEB #M0371S). The digested library was ligated into pLibacceptorV2 using T7 DNA Ligase (NEB #M0318S), cloned into 5-alpha Electrocompetent *E. coli* (NEB #C2989K), and plated on LB + kanamycin (25 ug/mL) yielding approximately 2.3 million colonies estimated by counting simultaneously plated dilutions. After allowing for 24 hours of growth on plates, the library was scraped and resuspended in LB, and then 800 million cells (based on OD_600_) were inoculated in 450 mL LB + kanamycin (25 ug/mL) overnight. Unless stated otherwise, all plasmids were isolated using a Qiagen Plasmid Plus Maxiprep Kit (#12963) and concentrated using a Promega Wizard SV Gel and PCR Clean-up System (#A9281).

In order to clone the RiboJ::sfGFP reporter construct, the library was digested using NEB’s BsaI-HF and NheI-HF with the addition of rSAP. The reporter construct was digested using NEB’s BsaI-HF and NcoI-HF. Similarly to the previous cloning step, the reporter was cloned into the library using T7 DNA Ligase, cloned into 5-alpha electrocompetent *E. coli*, and plated on LB + kanamycin (25 ug/mL), yielding 6.8 million colonies. The completed plasmid library was isolated as stated above.

### Isolation of genomic fragment library

To isolate genomic fragments, 10 ug of *E. coli* MG1655 gDNA was sheared using a Covaris focused ultra-sonicator. The settings used were as follows: Duty factor was set to 10%, Intensity was set to 4, cycles/burst was set to 200, and time was 60 seconds. The sheared gDNA was ran on a 3% TAE agarose gel and fragments between 200 and 300 bp were extracted using a Zymoclean Gel DNA Recovery Kit and eluted in 18 uL water. All 18 uL of the extracted fragments were end repaired using Enzymatics End Repair Mix (Part # Y9140-LC-L) following manufacturers protocols, cleaned using 45 uL (1.8x volume) of Agencourt AMPure XP Beads (#A63880), and eluted in 20 uL of water. The 20 uL eluate was A-tailed following the New England Biolabs protocol:

Reaction:

uL End-repaired DNA
uL NEB Buffer 2 (10x)
uL dATP (10mM)
uL Klenow Fragment (3’ −> 5’ exo-) (Enzymatics #P7010-HC-L)
uL Nuclease-free water

The reaction was Incubated for 30 minutes at 37°C, then heat inactivated for 20 minutes at 75°C before cleaning using 90 uL Agencourt AMPure XP beads and eluting in 20 uL water. Y-adapters to facilitate fragment amplification and barcoding were ligated to the A-tailed fragments using the following reaction mix:

Reaction:

uL A-tailed DNA
uL NEB T4 DNA Ligase Buffer (10x) (NEB #B0202S)
uL Y-adapter GU Y-Frag (25 uM)
uL NEB T4 DNA Ligase (NEB #M0202T)
uL Nuclease-free water

This reaction was incubated for 20 minutes at 25°C, heat inactivated for 20 minutes at 65°C, and subsequently cleaned using 90 uL Agencourt AMPure XP beads and eluting in 12 uL nuclease-free water.

### Barcoding and cloning of genomic fragment library

To barcode the genomic fragments, 1 uL of the processed fragments was amplified for 13 cycles using GU72 and GU116. This product was then cleaned using a Zymo Clean and Concentrator Kit and eluted in 12 uL nuclease-free water. For barcoding, 1 ng of this eluate was amplified for 10 cycles using primers GU72 and GU73. Following cleaning using a Zymo Clean and Concentrator Kit (#D40140), the library was digested using NEB’s SbfI-HF and XhoI.

This library was cloned following the same protocols as the TSS library. The transformation of the barcoded library yielded approximately 3.3 million colonies and the transformation after addition of the RiboJ::sfGFP yielded approximately 1.25 million colonies.

### Genomic promoter tiling library design

We used a custom peak caller on the single-nucleotide resolution strand-specific expression pileup generated from our genomic fragment library to define “peaks” of promoter activity. Our peak calling method is simple and conservative, as we wanted to tile the most active regions and keep the library size reasonable. We defined a peak as a continuous region with expression above an empirically determined threshold. We considered a continuous range of thresholds and for each evaluated the percentage of active TSSs, from our previous library, contained in a peak and determined an expression level of 1.1 was sufficient and captured 90% of active TSSs (data not shown). We required that each peak be at least 60 bp, and merged adjacent peaks that were within 40 bp, yielding 1753 and 1724 peaks for the minus and plus strands, respectively. We tiled each peak by synthesizing 150 bp windows across the region, with no overlap between adjacent tiles, yielding 48,491 peak tiles. Additionally, we included 1000 randomly generated 150 bp sequences to test what fraction of random sequence can drive expression. We included the same set of positive and negative controls as described in the TSS library design.

### Genomic promoter tiling library barcoding and cloning

The active TSS mutagenesis library was synthesized by Agilent and delivered lyophilized as a 10 pmol pool. The library was resuspended in 100 uL of TE pH 8.0 and 1 uL was amplified for 10 cycles using GU120 and GU121. This product was then cleaned using a Zymo Clean and Concentrator Kit and eluted in 12 uL nuclease-free water. For barcoding, 1 ng of this eluate was amplified for 8 cycles using primers GU120 and GU122. Following cleaning using a Zymo Clean and Concentrator Kit (#D40140), the library was digested using NEB’s SbfI-HF and XhoI.

This library was cloned following the same protocols as the TSS library. The transformation of the barcoded library yielded approximately 1.5 million colonies and the transformation after addition of the RiboJ::sfGFP yielded approximately 5.2 million colonies.

### Active TSS mutagenesis design

We systematically mutagenized all active TSSs from our initial TSS library to design a follow-up library. We used 500 negative controls to classify the TSS library into active and inactive TSSs. We set the active threshold at two standard deviations above the median expression for the negative controls, resulting in 2,670 active TSSs. We mutagenized the active sequence by scrambling 10 bp windows, sliding across the 150 bp at 5 bp intervals, resulting in 5 bp of overlap between adjacent scrambles. We scrambled the sequence using the existing 10 bp to preserve nucleotide content and selected the scramble that was most dissimilar to the original sequence out of 100 scrambling attempts. Our final library included 59,653 scrambled sequences and 2,057 unscrambled sequences. We also included the same set of negative and positive controls as described above for the TSS library, for a total library size of 62,322.

### Active TSS mutagenesis library barcoding

The active TSS mutagenesis library was synthesized by Agilent and delivered lyophilized as a 10 pmol pool. The library was resuspended in 100 uL of TE pH 8.0 and 1 uL was amplified for 12 cycles using GU123 and GU124. This product was then cleaned using a Zymo Clean and Concentrator Kit and eluted in 12 uL nuclease-free water. For barcoding, 1 ng of this eluate was amplified for 10 cycles using primers GU123 and GU125. Following cleaning using a Zymo Clean and Concentrator Kit (#D40140), the library was digested using NEB’s SbfI-HF and XhoI.

This library was cloned following the same protocols as the TSS library. The transformation of the barcoded library yielded approximately 3.7 million colonies and the transformation after addition of the RiboJ::sfGFP yielded approximately 5.2 million colonies.

### Library Barcode mapping

We used PCR to individually barcode each library sequence to quantitatively measure expression in our MPRA. Prior to genome integration, DNA-sequencing was performed to computationally map barcodes to sequences. A custom barcode mapper developed by Nathan Lubock ^97^ was used to collapse reads into a barcode-sequence map. We used two filtering steps for barcode quality. First, we required a minimum number of reads for every barcode, assuming reads that appear once or twice correspond to sequencing errors. Second, BBMap ^98^ was used to align the reads associated with a given barcode, and discarded barcodes that map to sequences that are too dissimilar to one another. A Levenshtein distance of 30 was used to discard barcodes that map to two very distinct sequences, while still allowing for a small number of sequence errors.

### Library integration into specific genomic loci

Library integration was performed as previously described ^38^.

The isolated plasmid library was digested with SalI-HF and NheI-HF to eliminate incompletely cloned plasmid before transformation into electrocompetent MG1655 with a landing pad engineered in the nth-ydgR locus and plating on LB + kanamycin (25 ug/mL). Colonies were resuspended in LB and 800 million cells were inoculated into 250 mL LB + kanamycin (25 ug/mL) and grown overnight. Several 2 mL frozen aliquots were made of this overnight culture.

The library was integrated into the *nth-ydgR* locus as follows. A frozen aliquot of MG1655 with a landing pad engineered in the reverse orientation at the *nth-ydgR* intergenic locus was transformed with the library and grown overnight in 200 mL LB + kanamycin (25 ug/mL). Following overnight growth, 400 million cells of this culture were seeded into 250 mL LB + kanamycin (25 ug/mL) + 0.2% arabinose (g/mL) and grown for 24 hours. After integration of the library, the plasmid backbone was removed through heat-curing. From the 24 hour induced culture, 800 million cells were inoculated into 80 mL of LB + kanamycin (25 ug/mL) and grown at 42 °C for approximately 1.5 hours before reaching an OD 600 = 0.3. Upon reaching exponential growth, 200 million cells from this culture library were plated and grown for 16 hours at 42 °C. Heat-cured plates were scraped and resuspended in LB and 400 million cells were inoculated into 200 mL LB + kanamycin (25 ug/mL). This culture, consisting of our integrated and heat-cured library, was grown overnight at 37 °C and several frozen 2 mL aliquots were made.

To test the TSS library in the *essQ-cspB* and *ybbD-ylbG* midreplichore regions, the same protocol was followed using strains engineered with landing pads in these intergenic regions.

### Library growth and harvest for expression measurements

To measure expression of all promoter libraries, libraries were grown and harvested as previously described ^38^ with minor changes to culture conditions.

For each library and biological replicates, a 2 mL frozen aliquot of the library was inoculated in 200 mL LB (BD#244620) with 25 ug/mL of kanamycin and grown at 30 °C overnight. The overnight cultures were used to seed new cultures at OD_600_ = .0005 and grown for approximately 5.5 hours at 30 °C until reaching an OD_600_ between = 0.5 and 0.55. The genomic fragment library was also grown in Minimal Media (Fisher Scientific #DF0485-17) with 0.2% glucose (g/mL) and 25 ug/mL of kanamycin for 10 hours at 30 °C until reaching an OD_600_ between = 0.5 and 0.55. Cultures were rapidly cooled to 0 °C in an ice slurry for two minutes. Three 50 mL aliquots were pelleted at 4 °C by centrifugation at 13,000xg for two minutes and the supernatants were poured out before snap-freezing the pellets in liquid nitrogen. Three 5 mL aliquots of each library were harvested using the same approach to be processed for genomic DNA extractions.

### RNA and DNA sequencing library preparation

RNA was extracted from 50 mL library pellets using a Qiagen RNEasy Midi kit (#75142) and 45 ug of each extract was concentrated using a Qiagen Minelute Cleanup Kit (#74204). Barcoded cDNA was generated from 25 ug of each concentrated RNA extract using Thermo Fisher SuperScript IV (#18090010) primed with GU101. The manufacturer’s protocol was followed aside from extending the reaction time to 1 hour at 52 °C. The cDNA reaction was cleaned using a Zymo Research DNA Clean and Concentrator kit (#D40140) before amplification. Barcoded cDNA was amplified via PCR for 13 cycles using primers GU59 and GU102. This reaction was cleaned using a Zymo Research DNA Clean and Concentrator Kit and 1 uL of this reaction was used in a second PCR for indexing and addition of flow cell adapters. The second PCR was for 8 cycles and utilized primers GU102 with either GU61, GU62, GU63, or GU64 (which add separate 6 bp indices).

gDNA was extracted from 5 mL cell library pellets using a Qiagen Gentra Puregene kit (#158567). Barcoded DNA was amplified from 1 ug of gDNA via PCR for 12-15 cycles using primers GU59 and GU60. The reaction was subsequently cleaned using a Zymo Research DNA Clean and Concentrator kit. To add sequencing adapters and indices to the library, 1 ng of this reaction was subject to a second PCR for 8 cycles using primers GU70 with either GU63, GU64, GU65, or GU66 (which add separate 6 bp indices). RNA and DNA sequencing libraries were cleaned using a Zymo Research Clean and Concentrator Kit (#D40140) before quantification using an Agilent Tapestation.

For each library, eight separate sequencing libraries were prepared: Four sequencing libraries for each RNA/DNA with two biological replicates and two technical replicates of each biological replicate. Biological replicates originated from separately grown and harvested glycerol stocks of each library. For each biological replicate, two RNA/gDNA extractions and sequencing library preparations (technical replicates) were performed in parallel. Libraries were submitted to the Broad Stem Cell Research Center at UCLA for sequencing on a HiSeq2500 or to the UCLA Translational Pathology Core Laboratory for sequencing on a NextSeq500. Raw sequencing data and promoter expression measurements are available on NCBI’s Gene Expression Omnibus (Accession no. GSE144621).

### RNA-Seq of MG1655 in M9 minimal Media and Rich LB media

To compare the promoter landscape to local transcriptional levels, RNA-Seq was performed on MG1655 grown in M9 minimal media (BD Difco #248510) supplemented with 0.2% glucose, 2 mM magnesium sulfate, and 0.1 mM calcium chloride. Similarly, RNA-Seq was performed for MG1655 grown in LB (BD#244620). Cells growth and RNA preps were prepared as previously described (see methods section titled: library growth and harvest for expression measurements). Samples were prepared using an Illumina TruSeq® Stranded mRNA Library Prep (#20020594) following manufacturers protocols to achieve strand-specific coverage. We note that no rRNA depletion was performed to preserve the fully intact transcriptional landscape. Samples were submitted to the UCLA TCGB sequencing core and sequenced on a Hiseq 4000.

### Standardizing Promoter Expression Quantification and Activity Thresholding

We processed the TSS, scramble, and peak tiling libraries using the same computational pipeline to facilitate comparisons between libraries. First, we use a set of 96 short synthetic positive controls, designed to span a range of activity^40,96^, to fit a robust linear regression (rlm function from MASS package) with the TSS library as the reference. Each library is standardized independently to the TSS library using the set of positive controls present in both libraries. Next, for each library we independently determined the level of background noise based on the median of 500 negative controls and subtracted this background from the newly fitted measurements. These steps standardize our data so we can train jointly across all datasets.

### -10 Motif and −35 Motif characterization

A position weight matrix from bTSSfinder was used to identify and score the best match to the −10 and −35 motifs within active tss-associated promoters, inactive tss-associated promoters, and a set of 500 negative controls. Best scores were reported regardless of position within the sequence. For all pairwise comparisons of active tss-associated promoters, inactive tss-associated promoters, and the negative controls, the distributions of motif scores were compared and a student’s t-test was performed to determine significance.

### Genomic fragment processing, alignment and promoter landscape quantification

To calculate fragment expression, we used measurements from DNA-seq and RNA-seq and excluded fragments with low expression (< 0.1) or high variance (5-fold difference in relative expression between biological replicates). To identify the coordinates of genomic fragments assayed using the MPRA, fragment sequences were aligned using bowtie2^99^ (version 2.3.4.3). To determine nucleotide-resolution calculations for promoter activity, we utilize the script, frag_expression_pileup.py. This script outputs WIG files in a strand-specific manner detailing the median expression of fragments overlapping each nucleotide position.

### Identification of minimal promoter regions

To identify minimal sequences necessary for promoter activity, contiguous stretches of candidate promoter region peak tiles were grouped and the minimal shared overlapping region was identified. Peak tiles above the expression threshold were identified and grouped together if they shared an overlap of at least 110 bp of their 150 bp total length. The minimal region necessary for promoter activity was found by determining the overlap of the outermost sequences within a contiguous stretch of tiles.

### Determining promoter-gene associations

To assign genomic promoter peaks to their regulated genes, peaks were first assigned specific nucleotide positions by identifying the maximum activity score within a peak. Promoter peaks were considered intragenic if their maximum scoring nucleotide overlapped with a gene coordinate. For peaks whose maximum scoring nucleotides were within intergenic regions, regulated genes were assigned by identifying the first downstream gene within 500 bp. Once gene associations were identified, promoter peaks were labeled sense or antisense depending on whether the regulated gene shared strand orientation with the promoter peak

### RNA-Seq alignment and genome transcript coverage

RNA-Seq analysis was performed using the script *RNAseq_LB_processing.sh* or *RNAseq_M9_processing.sh*. This script trims reads using the trimmomatic software (ver. 0.36+dfsg-3) and aligned to the MG1655 reference genome (U00096.2) using Hisat2 ^100^ (ver. 2.1.0-1). Genome nucleotide-resolution coverage was determined using Samtools depth (ver. 1.7-1). Meta-analysis across gene groups (as in figure 3B), was performed using Deeptools^101^ (ver. 2.5.6). Gene expression coverage (as in figure 4B) was calculated using custom script *wig_over_bed.py*, which calculates the total (.wig) coverage across reported *E. coli* genes. In all cases, default parameters were used with the exception of allowing for strand-specific quantifications.

### Amino acid and codon bias within intragenic promoters

Amino and codon usage was characterized within intragenic promoters and compared to all *E. coli* coding regions. To identify intragenic promoters, minimal regions necessary for promoter activity were identified by cross referencing genomic coordinates to reported genes. Reported gene coordinates were acquired from RegulonDB Version 8.0^83^. Once intragenic promoters were identified, nucleotide triplets were extracted while conserving the reading frame of the overlapping gene. Similarly, nucleotide triplets were extracted from all reported *E. coli* coding regions after filtering out sequences which did not have nucleotide lengths of a multiple of three. For these extracted sequences, codon frequencies were normalized to their relative abundance compared to other codons encoding the same amino acid. Amino acid frequencies were normalized to the total number of amino acids within each group. Significantly enriched or depleted codons were identified by performing a chi-squared test within each amino acid group and adjusting the p-value using FDR. Significantly enriched or depleted amino acids were identified by performing a chi-squared test for each amino acid relative to the total pool of amino acids and adjusting the p-value using FDR.

### Comparison of condition-dependent promoter and gene activation between rich and minimal media

To identify condition specific promoters, coordinates of candidate promoter regions identified in both M9 and LB conditions were compared to identify overlaps. Coordinates of promoter peaks were cross compared between conditions using the bedtools intersect tool (bedtools v2.27.1) and considered unique to a particular condition if they had no overlap between conditions. To identify regions that were activated between conditions, we compared the relative promoter activity between conditions at all positions in the genome and identified stretches greater than 60 bp that exhibited over 2-fold difference in activity. Regions were called using custom script run_differential_wig.sh available on the Github repository. To identify genes being expressed by differentially active regions, intergenic differentially active regions and matched these to the nearest downstream gene within 500 bp.

### Identification of SEED subsystem annotations enriched in differentially activated genes

To identify genetic functions associated with condition-dependent genes, the *E. coli* MG1655 K-12 genome (Genbank: U00096.2) was annotated using the SEED and RAST webserver ^67,68^. Genes within 500 bp downstream of promoter regions activated by condition were identified and associated with activation in LB or minimal media. For each media condition, genes were grouped by functional categories and the number of genes for each category was tallied.

### Identification of condition dependent TFBSs

The TFBS content of promoter peaks unique to each condition was evaluated by cross-referencing with TFBSs reported by RegulonDB ^83^ (Release 8.8). Genomic regions activated in each condition were assigned TFBSs based on overlapping genomic coordinates using the bedtools intersect tool (bedtools v2.27.1) with default parameters and ignoring strand assignments. Incidents of each TFBS overlap were quantified between conditions and normalized to incidents per 100,000 bp of promoter peak sequence.

### Identification of statistically significant scrambling promoter variants

We identified scrambling promoter variants that significantly altered expression compared to the wild-type (WT) variant in the script scramble_ttest.Rmd. We considered each scramble and barcode combination as an independent observation, rather than summarizing expression as an average across all barcodes. A two-sample two-sided Student’s t-test (t.test) was performed to test for a significant difference in mean expression levels between barcodes for a scrambled variant and barcodes for the corresponding WT variant. We performed multiple testing correction and identified 1,885 scrambles that increase expression and 5,408 that decrease expression relative to the WT variant, at a false discovery rate of 1%.

Next, bedtools merge was used to merge overlapping adjacent scramble variants to produce “merged” scrambles. These merged sites correspond to a continuous scrambled region that induced significant changes in expression. We identified 1,414 merged scrambles that increased expression and 1,903 merged scrambles that decreased expression, and scrambles were merged separately based on effect.

### Comparison of identified regulatory regions to RegulonDB annotations

We compared our identified merged scramble sites to existing RegulonDB annotations. We used bedtools intersect and required that 10% of the TFBS overlapped with a merged scramble site to count as an overlap. Next, we assessed whether the expression effect seen in our MPRA agreed with the direction of effect of the TFBS as indicated in RegulonDB. A merged scramble site was marked as “concordant” if any of the component scrambles agreed with existing annotation, and not concordant otherwise.

### Machine learning models

We implemented several machine learning models, independently trained for both classification and regression. All reproducible code is provided in the Github (https://github.com/KosuriLab/ecoli_promoter_mpra.git) and we will briefly describe each model and the appropriate parameters or implementation details.

#### Data processing

We standardized all datasets as detailed above in “Universal Promoter Expression Quantification and Activity Thresholding”. Next, we split our data, using custom scripts, into 75%/25% for training/testing based on genomic location, ensuring the splits are equidistant from the origin, to avoid overfitting (define_genome_splits.py). Briefly, we split the genome into eight chunks, with the first and last chunk adjacent to the origin of replication. We designated the second and seventh chunk as the test set and remaining chunks as training set. This splitting maintains roughly the same distance from the origin between the training and test sets to avoid any potential effects of genome location. Many of our library designs include high overlap between adjacent positions in the genome. Splitting by genome location mitigates inflated performance due to highly similar sequences present in both train and test sets. Across the three libraries (TSS, peak tiling, scramble) there are 87,164 training samples and 30,392 test samples.

We trained models for both regression and classification. Our data was skewed toward negative examples, with many samples near our determined threshold. For classification, we created a buffer around the threshold and only include sequences with expression <= 0.75 as negatives and >= 1.25 as positives and labeled sequences as active or inactive. Our training set was reduced to 53,326 samples and testing set to 18,567 samples.

We used the classification models to predict probabilities, instead of the class, to derive precision-recall curves.

#### Simple model with promoter features

For the models in this section we created features only for the TSS library because it is closest to endogenous sequence and is a smaller dataset. The training and test sets were split by genomic location, as described above, with 13,118 training samples and 4549 testing samples.

We created a simple model which incorporates four features related to promoter function. We calculated the maximum position weight matrix (PWM) score using motifs from bTSSfinder^102^ for both the −10 and −35 core promoter motifs. We scanned the −10 and −35 PWM individually and took the max score at any position using scoring functions from the Bioconductor package Biostrings^103^. Next, we scanned the sequence with −10 and −35 PWM jointly, allowing either 16, 17, or 18bp spacing in between the PWMs, reflecting common spacer lengths between core motifs. We assigned the “paired” max score as the max score at any position in the sequence across the three length options. Finally, we calculated the GC content (percentage) as this has been shown to be negatively correlated with promoter strength^104^. We constructed models in R with these four features and fit 1) a linear regression (lm), 2) a linear regression on the log-transformed expression values (lm), and 3) a logistic regression (glm, family = ‘binomial’, type = ‘response’).

We trained the gapped k-mer SVM (gkm-SVM^105^) model on only the TSS dataset because the model is suited for training sets < 20,000. The training and test sets were split by genome position as described above. We specified a word length = 10 with 8 informative columns (L = 10, K = 8).

#### K-mer frequencies and simple models (linear regression, logistic regression, partial least squares regression, partial least squares discriminant analysis)

All of the models described in the remaining sections were trained using all three combined datasets, as described above.

We created a feature set based on k-mer frequencies, with k-mers ranging in length from 3 to 6-mers. We generated feature sets and trained models in python. For simpler models we performed an additional feature selection step using custom scripts (kmer_feature_generator.py).

We trained four models:

● linear regression (statsmodel.api.OLS)
● logistic regression (sklearn.linear_model.LogisticRegression())
● partial least squares regression (sklearn.cross_decomposition.PLSRegression())
● partial least squares discriminant analysis (sklearn.cross_decomposition.PLSRegression() on binary dependent variable)

For each k-mer, we computed the frequency in a set of random genomic sequences, the same length and size of the training set. We include a k-mer if the absolute correlation with expression is greater than the “random” k-mer frequency, resulting in 4800/5440 filtered k-mers. We chose partial least squares regression because it projects the input features onto a new space and is better equipped to handle a large number of features with high collinearity.

#### Random forest regression and classification

Next, we trained a random forest, for both regression (sklearn.ensemble.RandomForestRegressor()) and classification (sklearn.ensemble.RandomForestClassifier()). We train on one-hot encoded DNA as a comparison to the neural network model, although random forest is not well suited to categorical input features. To compensate for this, we trained the random forest using frequencies of all 6-mers and observed improved performance.

#### Multi-layer perceptron and neural networks

We trained a multi-layer perceptron for both regression (sklearn.neural_network.MLPRegressor()) and classification (sklearn.neural_network.MLPClassifier()). MLPs are a class of feedforward artificial networks and are “vanilla” neural networks consisting of an input layer, hidden layer, and output layer. We used two different feature sets: frequency of all 3-to 6-mers and frequency of only 6-mers. Feature sets were standardized with sklearn.preprocessing.StandardScaler() to remove mean and scale to unit variance. We trained all four models with the following parameters: alpha = 0.005, hidden_layer_sizes=(800, 30), solver = ‘lbfgs’, random_state=1, max_iter=10000, early_stopping=True, learning_rate=’adaptive’, tol=1e-8.

We trained a convolutional neural network (CNN) on one-hot encoded DNA sequence for both regression and classification. We performed hyperparameter tuning and training using ^84^, a toolkit for working with CNNs built on keras. We performed a random hyperparameter search for a three-layer CNN for 100 combinations and the optimal parameters are listed below.

Regression:

● Dropout: 0.1340735187802852
● Pooling width: 16
● Convolutional filter width (for each layer): 16, 17, 18
● Number of filters (for each layer): 19, 39, 54

Classification:

● Dropout: 0.45541334972592196
● Pooling width: 7
● Convolutional filter width (for each layer): 8, 29, 29
● Number of filters (for each layer): 99, 87, 60

