## Supplementary Figures for "Genome-wide Functional Characterization of Escherichia coli Promoters and Sequence Elements Encoding Their Regulation"

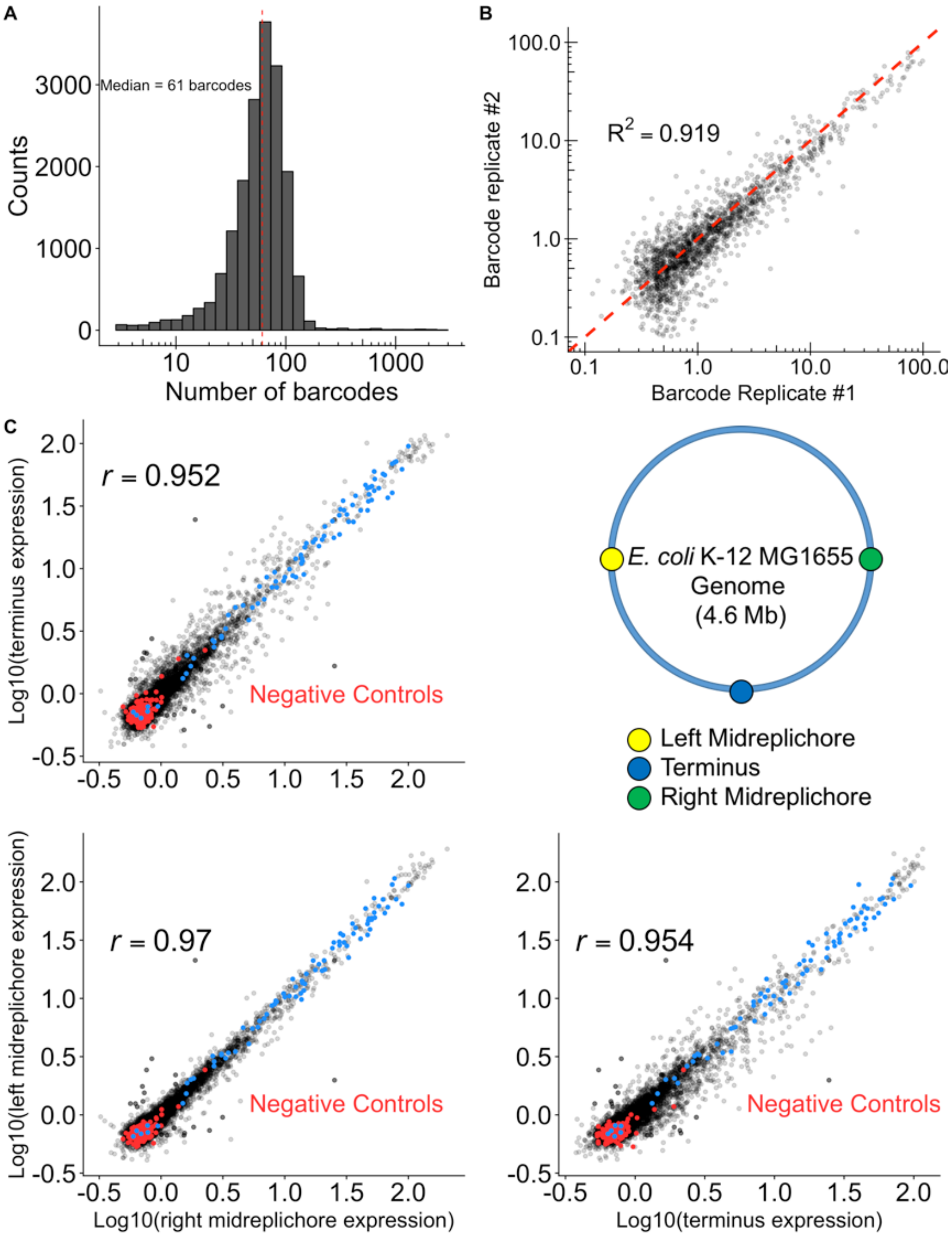


**Figure S1, related to Figure 1) TSS-associated promoters are represented by multiple barcodes and provide replicable measurements between genomic positions. A)** Distribution of the number of barcodes measured per TSS-associated promoter (Median = 61 barcodes). **B)** Comparison of separately barcoded experimental replicates for 1,824 TSS-associated promoters. Red line denotes y=x ($R^2$ = 0.919, $p < 2.2 \times 10^{-16}$). Barcode replicate #1 corresponds to promoters tested in the TSS variant library while Barcode replicate #2 corresponds to identical variants tested with the scanning mutagenesis library. **C)** Comparison of TSS-associated promoter measurements when integrated into distant regions of the *E. coli* chromosome ($p < 2.2 \times 10^{-16}$ for all correlations).

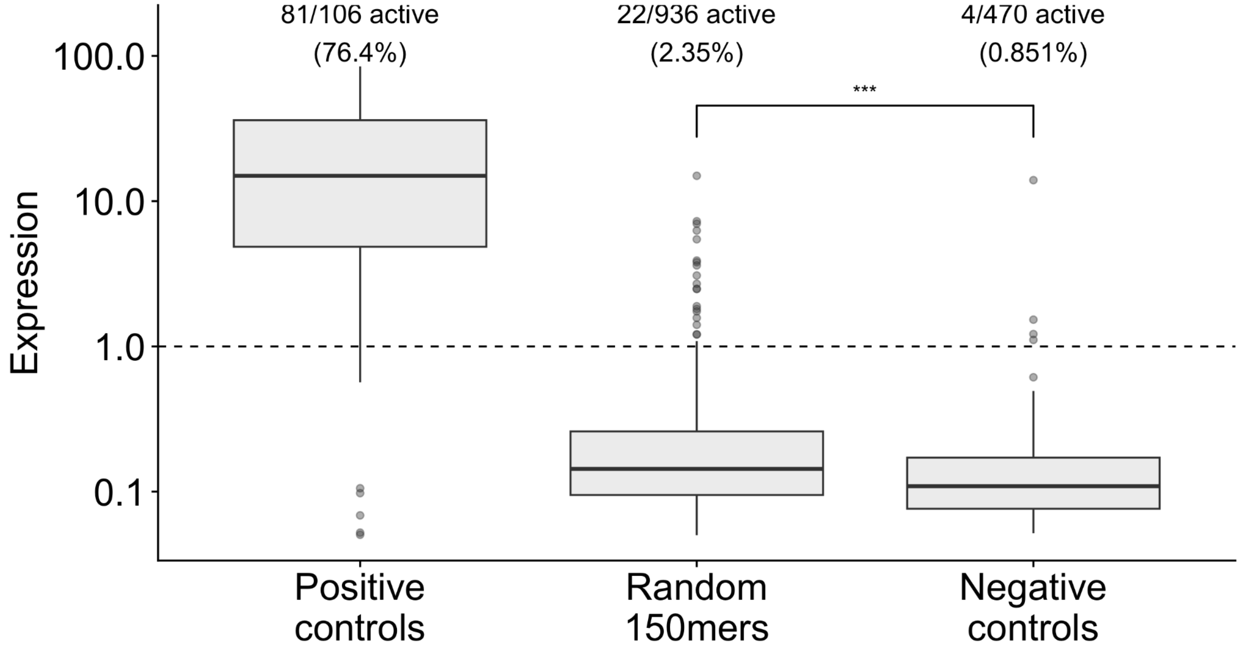


Figure S2) **An appreciable number of random 150mer oligos encode promoter activity.** Here we show the proportion of active sequences within each group of sequences tested using the promoter MPRA. Dashed line at 1 indicates the activity threshold as determined by the negative controls, which were measured in parallel with each group.

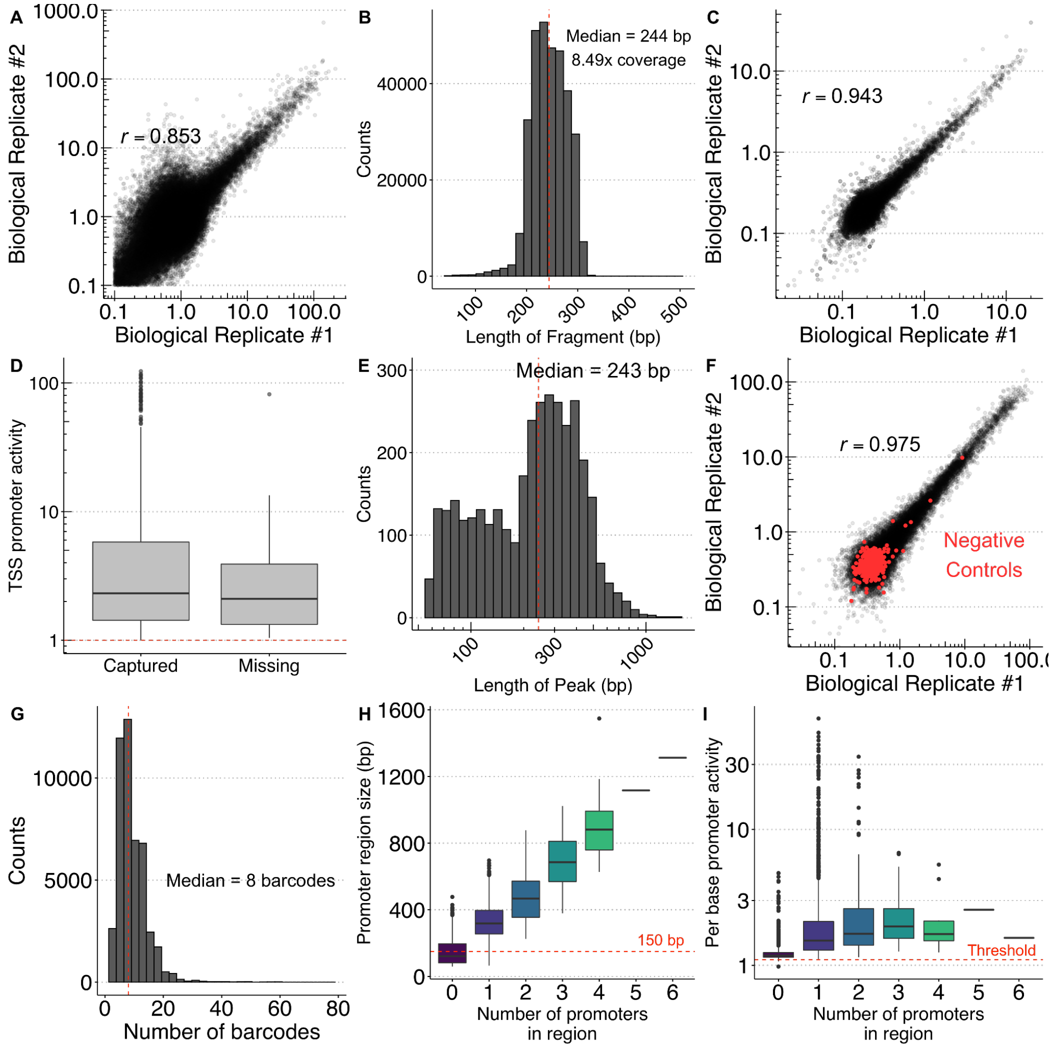


**Figure S3, related to Figure 2) Statistics of genomic screen for promoter activity in LB media. A)** Comparison of genomic fragment expression measurements between biological replicates. Genomic fragments with >5-fold difference in expression between biological replicates or expression under 0.1 were removed from downstream analysis ($r$ = 0.853, $p < 2.2 \times 10^{-16}$). **B)** Distribution of the lengths of genomic fragments assayed for promoter activity. **C)** Comparison of promoter activity measurements for 50,000 randomly sampled single-nucleotide positions between biological replicates ($r$ = 0.984, $p < 2.2\times10^{-16}$). Single-nucleotide position activity was determined by calculating the median expression of all fragments overlapping each individual position. **D)** Promoter activity measurements for active TSSs recaptured by genomic fragment screen compared (N=2,285) to TSSs that were not captured (N=386). **E)** Distribution of the lengths of genomic regions exhibiting promoter activity above our threshold. **F)** Comparison of peak tiling variant measurements between biological replicates ($r$ = 0.975, $p < 2.2\times10^{-16}$). **G)** Distribution of the number of barcodes per variant within the peak tiling library. **H)** Distribution of the size of promoter regions identified from the genomic fragment screen separated by their number of distinct minimal promoters. **I)** Distribution of the average per base promoter activity of promoter regions identified from the genomic fragment screen separated by their number of distinct minimal promoters.

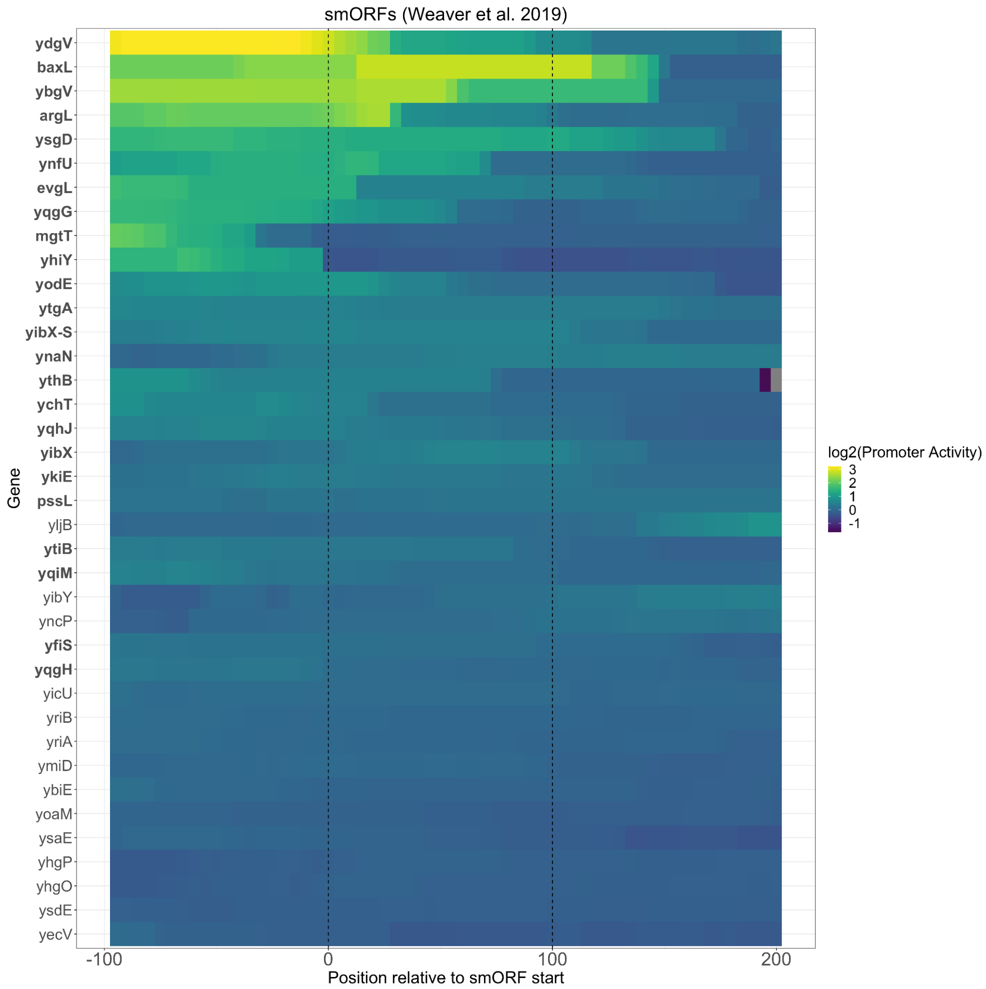


**Figure S4, Related to figure 2) Sense promoter activity upstream of 38 identified smORFs in LB media.** We detect promoter activity within 100 bp upstream of 24/38 smORFs. The names of smORFs with active promoter activity detected upstream are highlighted in bold. The lengths of all smORFs are scaled to 100 bp.

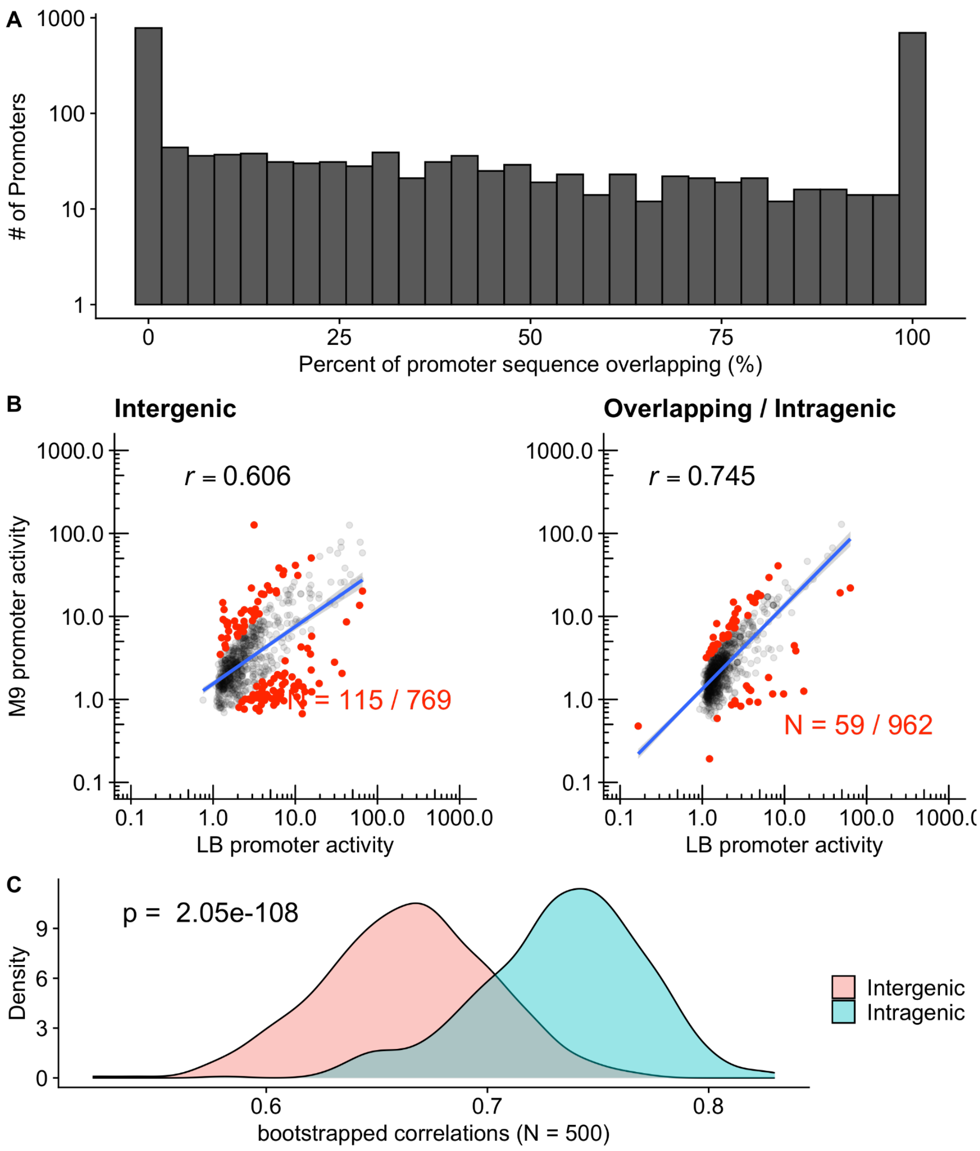


**Figure S5, related to Figure 3) Intragenic promoters exhibit reduced differential promoter activity in response to environmental conditions compared to intergenic promoters. A)** Distribution of the percentage of promoter sequence overlapping intragenic regions. Most promoter sequences were either fully intragenic or intergenic. **B)** Comparison of intergenic and intragenic promoter activity in LB vs M9 media for minimal promoter regions identified in tiling assay (**Figure 3A**). Intragenic promoters have more consistent activity between rich and M9 minimal media conditions. Red points highlight regions the most differentially active regions beyond the 95% confidence interval. **C)** Bootstrapped correlations between M9 and LB media for intragenic and intergenic promoters. The difference in correlations of promoter activity between conditions is statistically significant, indicating intergenic promoters exhibit greater condition-dependent activity ($p < 1 \times 10^{-16}$, Wilcoxon rank-sum test). 500 replicate bootstraps were performed.

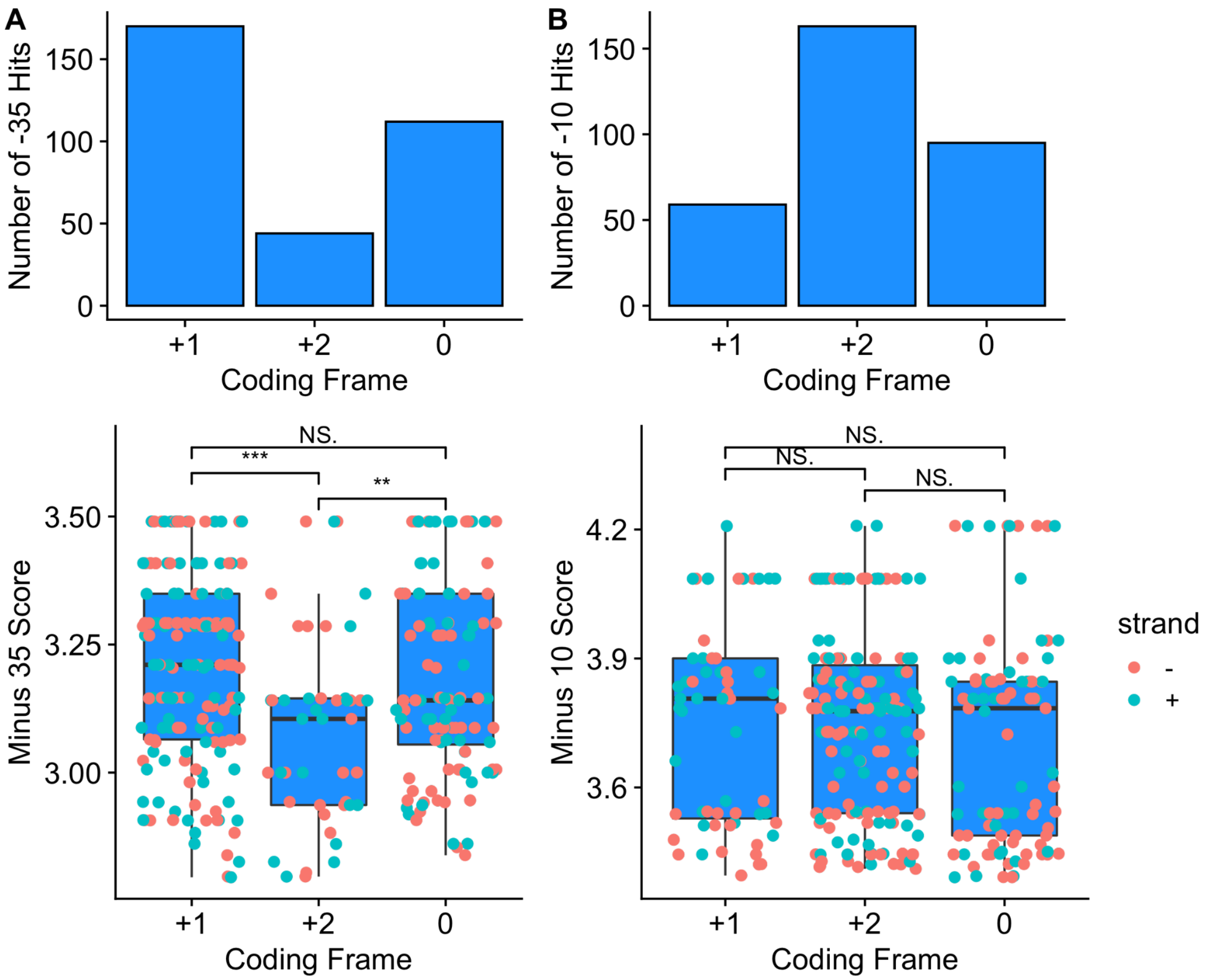


**Figure S6, Related to Figure 3) Intragenic σ70 promoter elements are depleted from specific reading frames within genes. A)** (Top) Number of -35 element hits within sense-intragenic promoters encoded within each reading frame. (Bottom) PWM scores of -35 elements within sense-intragenic promoters encoded within each reading frame. **B)** (Top) Number of -10 element hits within sense-intragenic promoters encoded within each reading frame. (Bottom) PWM scores of -10 elements within sense-intragenic promoters encoded within each reading frame.

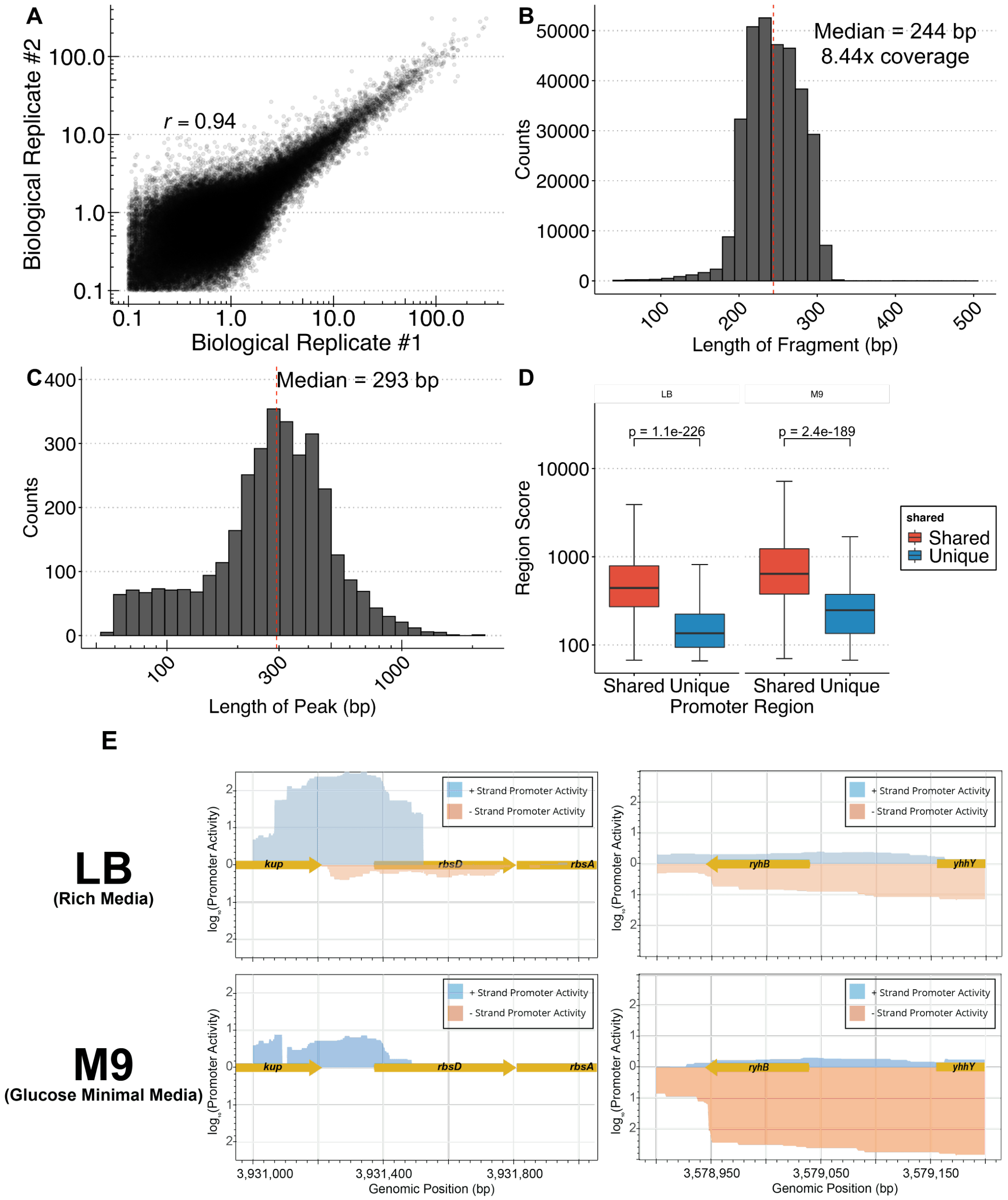


**Figure S7, related to Figure 4) Statistics of genomic fragmentation library in glucose minimal media. A)** Comparison of MPRA measurements between biological replicates in glucose minimal media. Genomic fragments with >5-fold difference in expression between biological replicates or expression under 0.1 were removed from downstream analysis ($r$ = 0.94, p < 2.2 x $10^{-16}$).**B)** Distribution of fragment lengths measured in glucose minimal media. **C)** Distribution of the sizes of promoter regions identified in glucose minimal media. **D)** Comparison of promoter region scores for regions that are unique to each condition or shared between conditions. **E) (Left)** *rbsDACBKR* operon promoter activity in LB media (top) and glucose minimal media (Bottom). **(Right)** *ryhB* promoter activity in LB media (TOP) and glucose minimal media (Bottom).

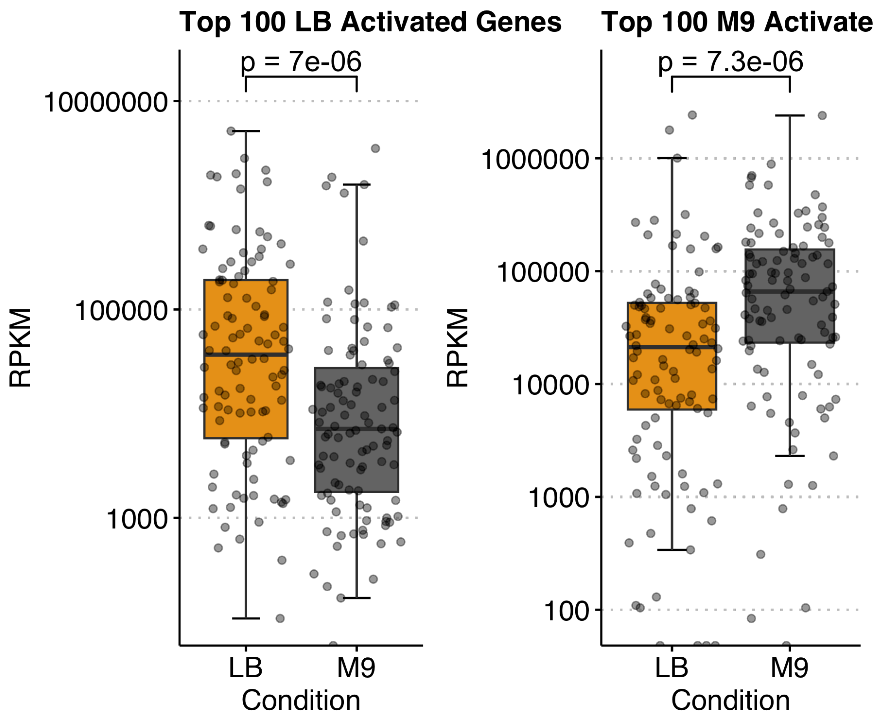


**Figure S8, related to Figure 4)** RNA-Seq expression of the 100 genes with the highest increase in promoter activity in (Left) LB media and (Right) glucose minimal media. Increase in the upstream promoter activity of these genes corresponds with significant transcriptional upregulation.

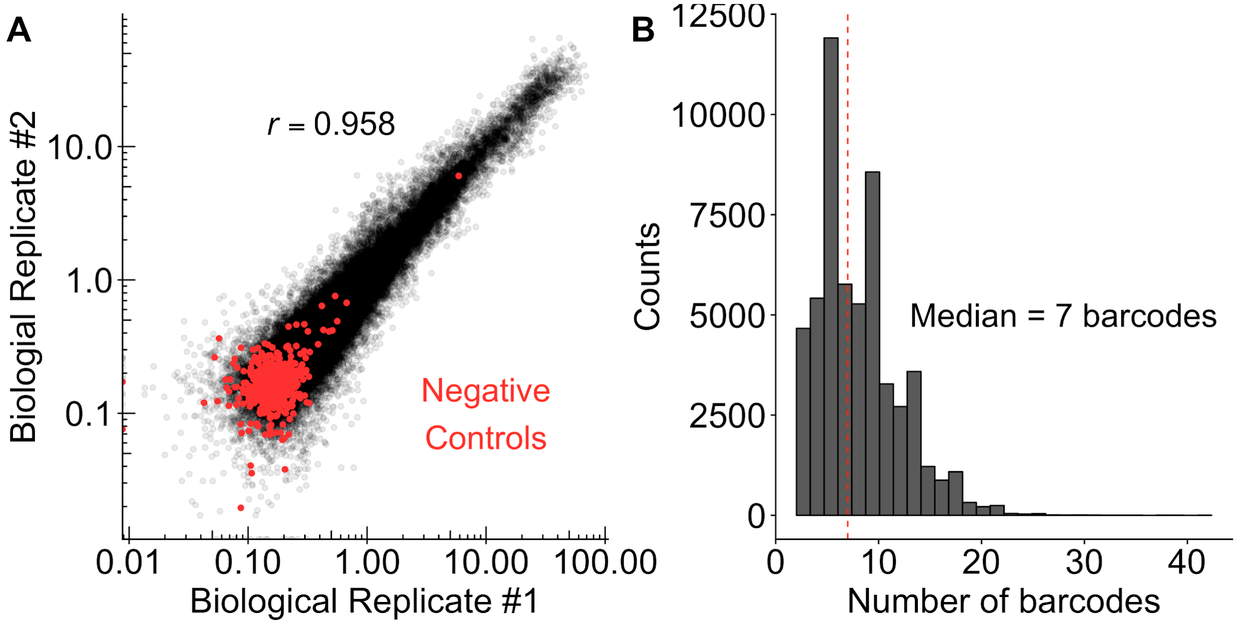


**Figure S9, related to Figure 5) Quality control for TSS scanning mutagenesis library. A)** Comparison of MPRA promoter activity measurements between biological replicates (*r* = 0.958, $p < 2.2 \times 10^{-16}$). **B)** Distribution of the number of barcodes per variant in TSS scanning mutagenesis library.

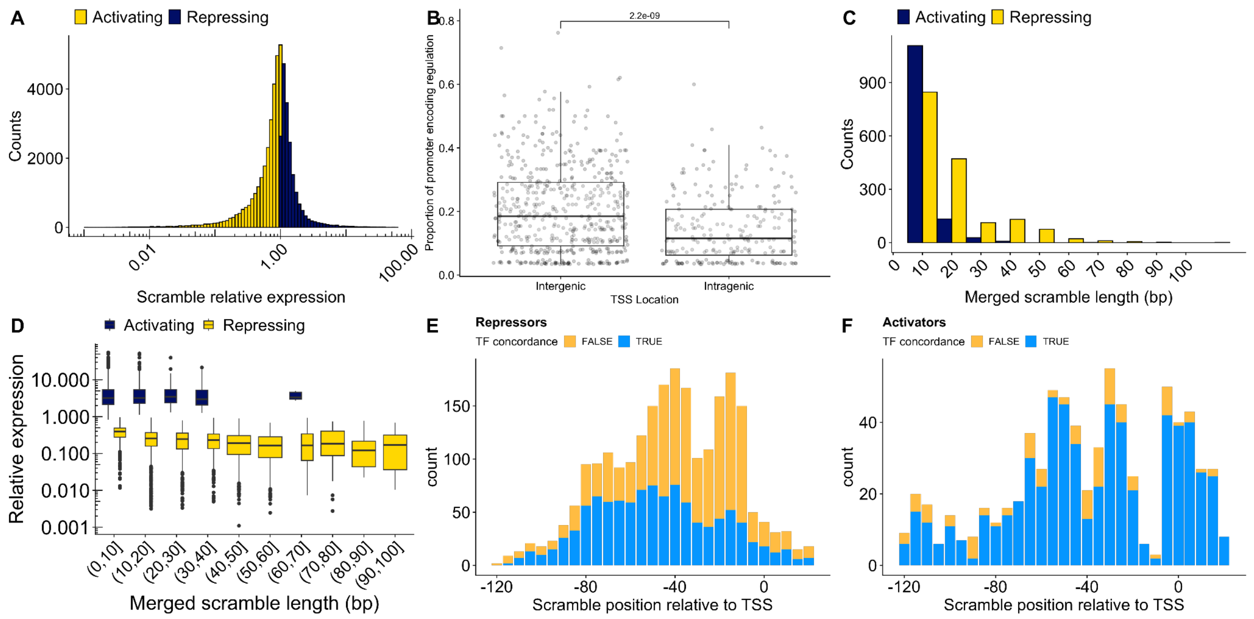


**Figure S10, related to Figure 6) Global identification of *E. coli* regulatory motifs by scanning mutagenesis. A)** Distribution of the effects of scrambling mutations regulatory regions on . **B**) A great proportion of intergenic promoter sequences impact expression when mutagenized compared to intragenic promoter sequences ($p$ = 2.2 x $10^{-9}$, Wilcoxon rank-sum test). **C)** Distribution of significant scramble lengths after merging contiguous regions. **D)** Relative change in expression from merged scrambles by length. **E,F)** Agreement between regulonDB TFBS annotations and effects of scrambling the overlapping region of the promoter for **E)** Repressors and **F)** Activators.
